# *Bacillus thuringiensis* Cry1A toxins divert progenitor cells toward enteroendocrine fate by decreasing cell adhesion with intestinal stem cells

**DOI:** 10.1101/2022.04.13.488147

**Authors:** Rouba Jneid, Rihab Loudhaief, Nathalie Zucchini-Pascal, Marie-Paule Nawrot-Esposito, Arnaud Fichant, Raphaël Rousset, Mathilde Bonis, Dani Osman, Armel Gallet

## Abstract

*Bacillus thuringiensis* subsp. *kurstaki* (*Btk*) is a strong pathogen toward lepidopteran larvae thanks to specific Cry toxins causing leaky gut phenotypes. Hence, *Btk* and its toxins are used worldwide as microbial insecticide and in genetically modified crops, respectively, to fight crop pests. However, *Btk* belongs to the *B. cereus* group, some strains of which are well known human opportunistic pathogens. Therefore, ingestion of *Btk* along with food may threaten organisms not susceptible to *Btk* infection. Here we show that Cry1A toxins induce enterocyte death and intestinal stem cell (ISC) proliferation in the midgut of *Drosophila melanogaster*, an organism non-susceptible to *Btk*. Surprisingly, a high proportion of the ISC daughter cells differentiate into enteroendocrine cells instead of their initial enterocyte destiny. We show that Cry1A toxins weaken the Cadherin-dependent adherens junction between the ISC and its immediate daughter progenitor, leading the latter to adopt an enteroendocrine fate. Hence, though not lethal to non-susceptible organisms, Cry toxins can interfere with conserved cell adhesion mechanisms, thereby disrupting intestinal homeostasis and enteroendocrine functions.

## Introduction

The gut lining is undergoing constant damage caused by environmental aggressors (pesticides, drugs, viruses, bacteria and toxins) ingested along with food. The gut quickly responds to these aggressions by accelerating its epithelium renewal to replace damaged cells. Over the past decade, studies in *Drosophila melanogaster* have contributed substantially to the understanding of the cellular and molecular mechanisms controlling the maintenance of intestinal homeostasis and regeneration. These mechanisms have proven to be highly conserved in the animal kingdom. In *Drosophila*, resident intestinal stem cells (ISCs) are the guarantors of this cell renewal process. Under normal conditions, asymmetric division of an ISC gives rise to a new ISC (to maintain the pool of ISC) and to a daughter progenitor cell that can commit to two different paths of differentiation. The enteroblasts (EBs) and enteroendocrine precursors (EEPs) are the precursors of enterocytes (ECs) and enteroendocrine cells (EECs), respectively (Guo et al., 2021; Joly and Rousset, 2020; Pasco et al., 2015). ECs are the main intestinal epithelial bricks constituting an efficient barrier against aggressors and are therefore their first victims. The damaged or dying ECs emit cytokines, which stimulate the proliferation of ISCs to augment the pool of EBs that will differentiate into ECs to replace the damaged ones (Bonfini et al., 2016; Osman et al., 2012). Two mechanisms underlying intestinal regeneration have been described. The first one is the “cell renewal” model, which occurs under weak aggression conditions that does not induce EC apoptosis. In this case, ISC proliferation is low and the neo-EBs differentiate into ECs. However, this provokes a transient excess of ECs due to the absence of prior EC death. The gut cell homeostasis is subsequently reestablished by the removal of old ECs (Loudhaief et al., 2017). The second mechanism, called “cell replenishment” or “regenerative cell death”, occurs after a strong aggression that induces massive EC apoptosis. In this case, a rapid ISC proliferation is followed by the differentiation of EBs into ECs to replace the dying ones (Loudhaief et al., 2017; Vriz et al., 2014) without producing supernumerary ECs.

*Bacillus thuringiensis* (*Bt*) bacteria are largely used as microbial insecticides to fight crop pests. *Bt* is a Gram-positive sporulating bacterium belonging to the *Bacillus cereus* (*Bc*) group (Ehling-Schulz et al., 2019). It was first identified and characterized for its specific entomopathogenic properties due to the presence of a crystal containing specific Cry protoxins, which are produced during the bacteria sporulation (Rabinovitch et al., 2017). Among all the subspecies of *Bt* inventoried (http://www.bgsc.org/), spores of *Bt* subsp. *Kurstaki* (*Btk*) are used to specifically kill lepidopteran larvae that threaten crops, through a cocktail of Cry toxins made of Cry1Aa, Cry1Ab, Cry1Ac, Cry2Aa and Cry2Ab (Caballero et al., 2020). Cry toxins sequentially bind to different receptors present in the midgut to exert their cytotoxicity. Among those receptors, the ones named Bt-R that belong to the Cadherin transmembrane cell adhesion molecules are primordial for the Cry1A holotype of toxins, allowing them to bind to enterocyte brush borders. The other receptors—such as Alkaline phosphatases, Aminopeptidases N, and ABC transporters— appear to account for the cytotoxicity that Cry exert toward susceptible organisms (Adang et al., 2014; Gao et al., 2019; Li et al., 2020; Liu et al., 2018b). In susceptible insects, upon ingestion of spores and crystals, the basic midgut pH dissolves the crystals, releasing the Cry protoxins. Then, digestive enzymes cleave Cry protoxins (130kD and 72kD for proCry1A and proCry2A, respectively) into activated Cry toxins (around 67kD) allowing them to bind their midgut receptors. Thereby, Cry toxins form pores in the plasma membrane of ECs, ultimately leading to their death. An alternative mode of action of Cry toxins suggested that Cry binding to Cadherin induces an intracellular flux of Mg^2+^ resulting in EC apoptosis (Castella et al., 2019; Mendoza-Almanza et al., 2020). In both models, toxin-induced breaches within the gut lining allow bacteria (spores and vegetative cells) to reach the internal body cavity, generating a septicemia and subsequent death of the lepidopteran larvae within 2 or 3 days after ingestion of *Btk* spores (Mendoza-Almanza et al., 2020). It is assumed that *Btk* do not harm the intestine of non-susceptible organisms because, first, the intestinal pH is not suitable for the solubilization of the crystal of protoxins and, second, the Cry toxin receptors are absent from their gut epithelium (Rubio-Infante and Moreno-Fierros, 2016).

However, recent studies provide evidence that *Btk* also exhibits some adverse effects on non-susceptible organisms including humans. Indeed, strains of the *B. cereus* group, to which *Bt* belongs, are well-known worldwide food-poisoning pathogens causing diarrheal-type illnesses due to the production of enterotoxins during the vegetative phase in the intestine (Ehling-Schulz et al., 2019; Glasset et al., 2016; Jovanovic et al., 2021). Recently, *Bt* has also been implicated in foodborne outbreak events and the strains identified were indistinguishable from the commercial ones (Biggel et al., 2021; Bonis et al., 2021; Johler et al., 2018). Furthermore, we have shown that *Btk* spores and toxins at concentrations close to those recovered on vegetables after spraying induce growth defects and developmental delay in *Drosophila* larvae (Nawrot-Esposito et al., 2020). Increasing spore and toxin doses ultimately led to larval lethality (Babin et al., 2020). Cry toxins produced by *Btk* are also used in genetically modified crops (GMCs) (ISAAA, 2017), and it has been reported that GMC-produced Cry1Ab toxin is found in agricultural water stream networks at abnormally high doses that may affect the survival rate of non-susceptible insects (Rosi-Marshall et al., 2007). In a similar vein, laboratory studies have demonstrated genotoxic activity of Cry1Aa, Cry1Ab, Cry1Ac and Cry2A in zebrafish rearing water (Grisolia et al., 2009). Based on all these data, our aim in this study was to decipher the interaction of *Btk* and its toxins with the intestinal epithelium using *Drosophila melanogaster*, an organism non-susceptible to *Btk* Cry toxin and a well-established model for studying host-pathogen interaction mechanisms.

Using environmental doses of spores and crystals of protoxins recovered on vegetables after treatment, we first showed that crystals of *Btk* Cry protoxins induced moderate enterocyte death that triggers a quick cell replenishment by favoring symmetric ISC divisions. We then demonstrated that the crystals diverted a higher number of progenitor cells from their initial EC fate toward an EEC fate, generating an excess of EECs. Importantly, this effect was due to a weakened cell-cell interaction between ISC mother cells and progenitor daughter cells. We were able to rescue the Crystal-dependent excess of EECs by specifically overexpressing the DE-Cadherin in ISC and progenitor daughters, reinforcing the strength of the adherens junction between these cells. Moreover, we found that among the five *Btk* Cry toxins, only the Cry1A holotype was able to induce this EEC excess. Unexpectedly, we observed that *Btk* crystals are processed in the midgut of adult *Drosophila* as they are in that of susceptible-organisms, releasing activated Cry1A toxins. Hence, since our data demonstrate that Cry1A toxins disrupt conserved cellular processes, many non-susceptible organisms may exhibit an excess of EECs and consequently a disruption of their enteroendocrine functions.

## Results

### Crystals of *Btk* Cry protoxins induce EC death and stimulate proliferation of intestinal stem cells

During sporulation, *Btk* produces a crystal of protoxins that is lethal to lepidopteran larvae once ingested by the insect. To study the *Btk* effects on the non-susceptible organism *Drosophila melanogaster*, we orally infected flies with the SA11 *Btk* strain (hereafter named *Btk*^*SA11*^), which is widely utilized in commercial microbial insecticides. A suspension of spores/crystals in water was deposited on the fly medium corresponding to 10^6^ CFU (Colony Forming Unit) of spores per female for 4 cm². The impact on the gut of the spores alone, or the toxins alone, was also monitored using a *Btk* strain devoid of protoxin crystals (*Btk*^Δ*Cry*^) or purified crystals, respectively (see Materials and Methods).

The *Gal4/UAS* binary expression control system (Brand and Perrimon, 1993) allowed us to monitor first the effect of the spores/crystals on EC apoptosis by expressing the Caspase 3 sensor (Casp::GFP; (Schott et al., 2017) under the control of the *myo1A-Gal4* EC driver (*myo1A>Casp::GFP*). With this transgenic combination, the GFP is detectable only when the Caspase 3 is activated in ECs. As a negative control, we fed flies with water alone. In all our experiments, we focused our observations on the posterior midgut (R4 region, https://flygut.epfl.ch/overview) (Buchon et al., 2013) because this region is known to show a high stem cell renewal activity (Marianes and Spradling, 2013) and exhibits the strongest phenotypes (see below). One day post-ingestion, *Btk*^*SA11*^ or purified crystals induced moderate apoptosis of the ECs compared to the control (Figure 1A). However, the treatment with spores alone devoid of protoxin crystals (*Btk*^Δ*Cry*^) did not induce EC death.

**Figure 1:**
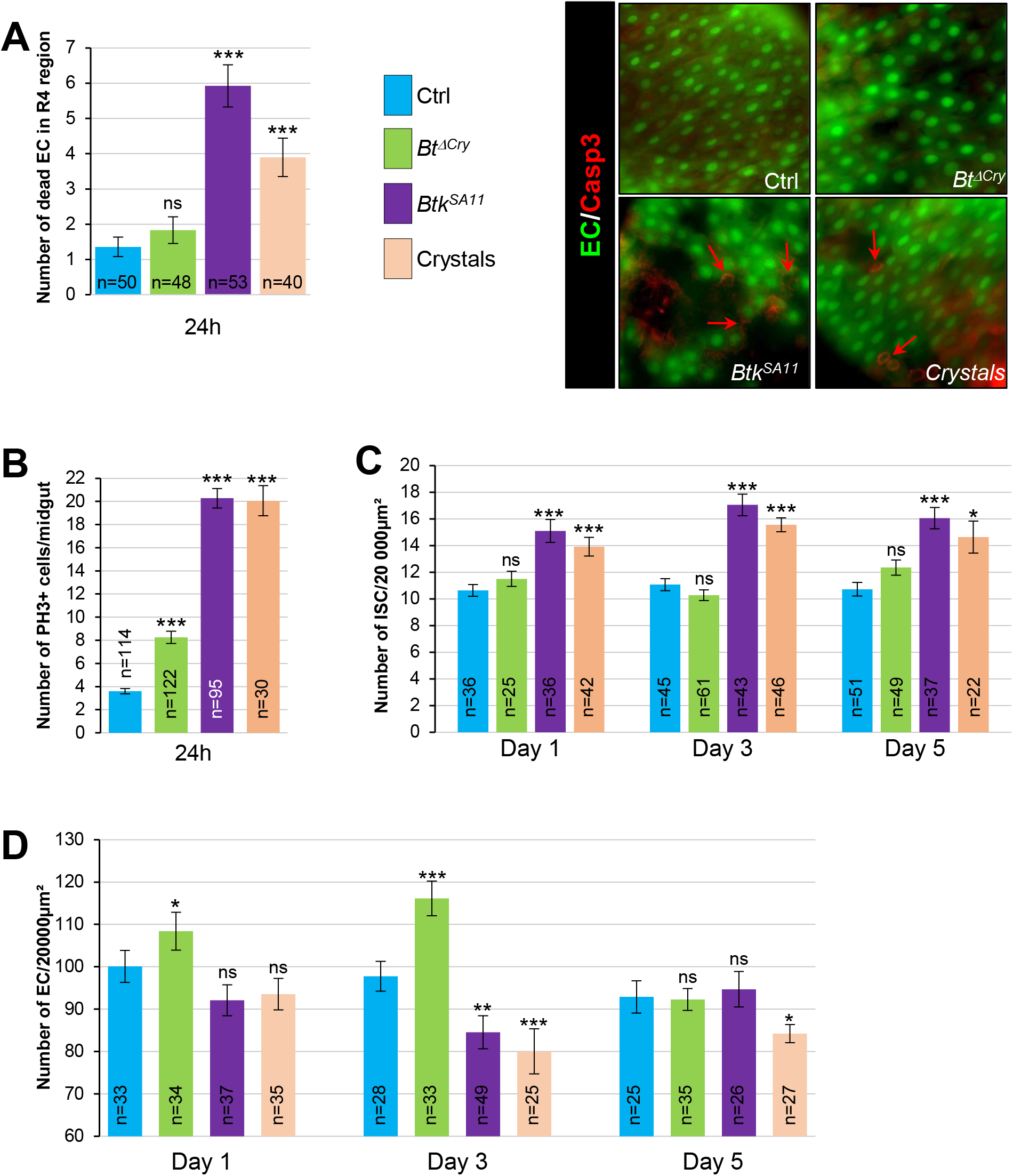
Crystals of *Btk* Cry protoxins induce EC death and stimulate proliferation of intestinal stem cells. (**A-D**) Flies were fed with water (blue, control-ctrl), *Btk*^*ΔCry*^ spores (green), *Btk*^*SA11*^ spores (purple) or Crystals (pink). The initial dose provided with the food to flies was 10^6^ CFU fly/4 cm² (this dose is used in all the following experiments presented in this article). **(A)** Left panel: Quantification of dead ECs 24h post ingestion (PI) in the posterior midgut (R4 region). EC apoptosis was monitored by expressing the Caspase 3 sensor (Casp:: GFP) using the *myo1A-GAL4* EC driver *(myo1A>Casp::GFP*). With this transgenic combination, the GFP is detectable only when the Caspase 3 is activated in ECs. Right panel: *myo1>GFP* midguts labeled with the anti-cleaved Caspase 3 antibody (red). **(B)** Quantification of mitoses using the anti-PH3 antibody in the whole midgut 24h PI. **(C)** ISC Number in the R4 region of *esg>GFP* flies 24, 72, and 120h PI. **(D)** EC number in the R4 region of *myo1A>GFP* flies 24, 72, and 120h PI. n=number of midguts; ns=not significant; * (P ≤ 0.05); ** (P ≤ 0.01), *** (P ≤ 0.001).

Induction of cell death is known to strongly induce ISC proliferation in the whole midgut (Biteau et al., 2008; Chatterjee and Ip, 2009; Jiang et al., 2009; Loudhaief et al., 2017). This prompted us to assess the number of ISC mitoses in the different conditions, using an Anti-phospho-Histone H3 antibody marking mitotic cells. As expected, ISC mitotic indexes were stronger upon oral infection with *Btk*^*SA11*^ spores or purified crystals than in the control (Figure 1B). In *Btk*^Δ*Cry*^ spore infection, mitotic figures were only moderately increased (Figure 1B). This is consistent with previous observations showing that a low dose (10^6^ CFU per *Drosophila*) of *Btk* vegetative cells (that do not produce and contain crystals of Cry protoxins) only moderately activates ISC proliferation without inducing EC apoptosis (Loudhaief et al., 2017). We confirmed this increase in ISC proliferation by counting the total number of ISCs in the posterior midgut (R4 region). To specifically mark ISCs, we expressed the GFP under the control of the *Dl-Gal4* driver that is expressed in ISCs and EEPs (Joly and Rousset, 2020), and we co-stained with an anti-Prospero (Pros), an EEP and EEC marker (ISCs were therefore GFP+, Pros-). While we observed an increase in ISC number with *Btk*^*SA11*^ spores or purified crystals, *Btk*^Δ*Cry*^ spores did not induce any increase (Figure 1C). This could be explained by the fact that the moderate stimulation of ISC proliferation by *Btk*^Δ*Cry*^ spores was not sufficient to promote a detectable increase in global ISC number. Another possibility could be that *Btk*^*ΔCry*^ spores favor asymmetric ISC division, giving rise to one ISC and one precursor (EB or EEP), while *Btk*^*SA11*^ spores or purified crystals promoted symmetric ISC division, giving rise to two ISCs. Indeed, numerous observations show that in the *Drosophila* midgut during intestinal homeostasis, the main mode of ISC division is asymmetric, accounting for about 70-80% of the mitosis. On the contrary, following an urgent need such as detrimental bacterial infection, ISCs divide symmetrically to allow faster regeneration of the tissue (de Navascues et al., 2012; Goulas et al., 2012; Joly and Rousset, 2020; O’Brien et al., 2011; Tian and Jiang, 2014; Zhai et al., 2017). To assess the mode of ISC division, we used the ReDDM *Drosophila* genetic tool (Antonello et al., 2015) under the control of the *Dl-Gal4* driver (*Dl*-ReDDM). This tool allows us to follow the progeny of the ISCs because they express a stable RFP (H2B::RFP) while the mother cells (the ISCs) express a labile GFP that disappears in less than 24 hours. After shifting the flies to 29°C to activate the *Dl*-ReDDM tool, the GFP was only expressed in Dl+ cells (i.e. the ISCs and EEPs) while the H2B::RFP was expressed in Dl+ cells but also stably transmitted to the progeny. When two cells in direct contact were ISC/ISC (GFP+/H2B::RFP+) or EEP/EEP (GFP+/H2B::RFP+/Pros+) or EB/EB (only RFP+), we considered they were coming from a symmetric division. In contrast, divisions were considered asymmetric when two neighboring cells displayed a different combination of immunolabelling, revealing that their identities were different. Interestingly while we observed a 33%/67% distribution of symmetric/asymmetric division in the control conditions (Figure S1A and S1B), the mode of division trended in favor of symmetric with *Btk*^Δ*Cry*^ spores (58.6%/41.4%) and was completely reversed with the *Btk*^*SA11*^ spores (74.2%/25.8%) (Figure S1A and S1B). Therefore, consistent with previous studies, the mode of ISC division appeared to reflect the degree of intestinal damage, the symmetric mode being preferentiality adopted when the damage was greater (i.e., when cell death occurred).

We next monitored the number of ECs using the fly strain *myo1A>GFP* allowing the expression of GFP in all ECs. *Btk*^*ΔCry*^ spores induced an excess of ECs at days 1 and 3 post-ingestion; the right number of ECs was recovered 5 days after ingestion (Figure 1D and Figure S1C). Along with the absence of EC apoptosis (Figure 1A) and the moderate induction of ISC proliferation (Figure 1B), our data strongly suggest that *Btk*^Δ*Cry*^ spores weakly damage the intestinal epithelium, reminiscent of the “cell renewal” process previously described after infection with poorly virulent bacteria (Loudhaief et al., 2017). On the contrary, ingestion of the *Btk*^*SA11*^ spores or purified crystals provoked a decrease in EC number 3 days post-ingestion (Figure 1D and Figure S1C) that we attributed to EC apoptosis (Figure 1A) (Loudhaief et al., 2017). A normal number of ECs was restored 5 days post-ingestion for *Btk*^*SA11*^ spores and to a lesser extent for purified crystals (Figure 1D and Figure S1C). Hence, *Btk*^*SA11*^ spores or purified crystals launch a process of regenerative cell death, inducing a strong proliferation of ISCs to quickly replenish the gut lining as previously described for strong pathogens (Vriz et al., 2014). Importantly, the ingestion of purified crystals containing the *Btk* Cry-protoxins recapitulates the midgut phenotypes caused by the ingestion of *Btk*^*SA11*^ spores.

### *Btk*^*SA11*^ spores induced an increase in EB, EEP and EEC numbers

During the course of our experiments we observed that many more EECs were apparently present in the flies that had ingested *Btk*^*SA11*^ spores compared with in the control flies (Figure S2A and S2C). As ECs derive from EBs and EECs from EEPs, we assessed the amount and the identity of the precursors in the different conditions of infection (H_2_O; *Btk*^*ΔCry*^ spores, *Btk*^*SA11*^ spores and purified crystals). To count the EBs, we used a *Gal4* strain of *Drosophila* driving GFP expression specifically in EBs (*Su(H)>CD8::GFP*). As expected, a significant increase in the number of EBs was observed between the first and the fifth day after ingestion of *Btk*^*ΔCry*^, *Btk*^*SA11*^ or purified crystals (Figure 2A). The EEP number was assessed using two markers: the GFP expressed in ISCs and progenitors (EBs and EEPs) using the *esg-Gal4* driver (*esg>GFP*) and a Pros staining which labels EEPs and EECs. Cells expressing both GFP and Pros corresponded to EEPs. While *Btk*^*ΔCry*^ spores did not modify the number of EEPs, ingestion of either *Btk*^*SA11*^ spores or purified crystals resulted in an increase in EEPs from day one onwards (Figure 2B and D). Since EEPs must differentiate into EECs, we then counted the differentiated EECs that were GFP-Pros+. No difference in EEC number was obtained with *Btk*^*ΔCry*^ spores compared to the control, whereas there was a net increase with the *Btk*^*SA11*^ spores and with the purified crystals (Figure 2C, D and S2A-D). Interestingly, though this event was rare, we also observed that EEPs could undergo a cycle of mitosis that might contribute to the increase in EEPs and EECs (Figure 2E) (Biteau and Jasper, 2014; Li et al., 2014; Zeng and Hou, 2015). Interestingly, we never observed such an increase in EEPs and EECs in the anterior part of the midgut (Figure S2E-I). Altogether, our results showed that the protoxin crystals of *Btk*^*SA11*^ were responsible for the excess of EECs in the posterior midgut.

**Figure 2:**
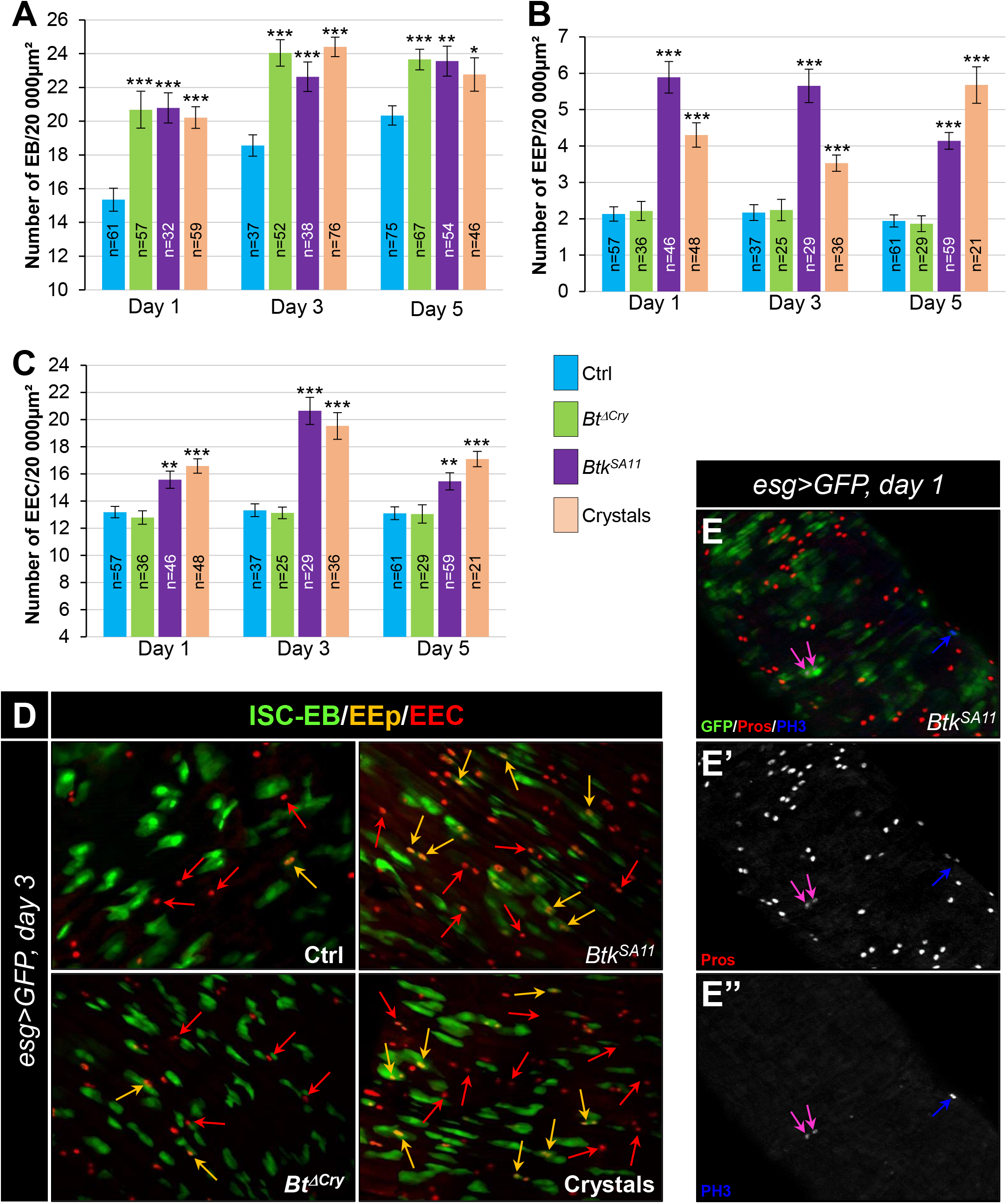
*Btk*^*SA11*^ spores induce an increase in EB, EEP and EEC numbers. (**A-D**) Flies were fed with water, *Btk*^*ΔCry*^ spores, *Btk*^*SA11*^ spores or Crystals. (**E-E”**) Flies were fed with *Btk*^*SA11*^ spores. (**A-C**) Control (water-ctrl): blue bars; *Btk*^*ΔCry*^ spores: green bars; *Btk*^*SA11*^ spores: purple bars; Crystals: pink bars. (**A**) Number of EBs in the R4 region of *Su(H)>CD8::GFP* flies 24, 48, and 72h PI. (**B** and **C**) Number of EEPs (B) and EECs (C) in the R4 region of *esg>GFP* flies 24, 48, and 72h PI. (**D-E”**) R4 region of *esg>GFP* flies labeled with anti-Pros (Red). GFP was expressed in ISCs, EBs and EEPs, and Pros was expressed in EEPs (yellow arrows in D) and EECs (red arrows in D). (E-E’’) PH3 staining (blue) marks mitosis. Pink arrows point to dividing EEPs and blue arrows point to dividing ISCs. n=number of samples; * (P ≤ 0.05); ** (P ≤ 0.01), *** (P ≤ 0.001).

### EEC excess arises from newborn EEPs after ingestion of crystals of *Btk*^*SA11*^ protoxins

To demonstrate that the excess of EEPs and EECs arose from proliferating ISCs caused by the ingestion of protoxin crystals, we used the ReDDM cell lineage tracing system using the *esg-Gal4* driver (*esg-ReDDM* flies). We chose to analyze the progeny at day 3 post-ingestion (Figure 3A), when the increase in EECs reached its peak (Figure 2C). According to the expression of specific cell markers and nucleus size, we could identify different cell types either that existed before the ingestion or that appeared after the ingestion of *Btk*^*ΔCry*^ spores, *Btk*^*SA11*^ spores or purified crystals. Identities of the different cell types were defined as follows: ISCs were GFP+ RFP+ DAPI+ with small nuclei; EBs were GFP+ RFP+ DAPI+ with bigger nuclei; EEPs were GFP+ RFP+ Pros+ DAPI+; new EECs were RFP+, Pros+ DAPI+; old EECs were Pros+ DAPI+, new ECs were RFP+ DAPI+ with big polyploid nuclei and old ECs were DAPI+ with very big nuclei (Figure 3 B-E”). In the control experiments, a few newborn ECs (red arrows in Figure 3B and B’, and Figure 3G) and rare newborn EECs (Figure 3F) appeared 3 days post-ingestion, reflecting the relative steady state of the cellular homeostasis. As expected for poorly virulent bacteria, ingestion of *Btk*^*ΔCry*^ spores induced the appearance of newborn ECs (red arrows in Figure 3C, C’ and Figure 3G) and only rare newborn EECs resulted (pink arrows in Figure 3C-C” and Figure 3E and G). Similarly, ingestion of *Btk*^*SA11*^ spores or purified crystals promoted the appearance of newborn ECs (red arrows in Figure 3D, D’, E and E’ and Figure 3G) but, strikingly, a high number of newborn EECs appeared (pink arrows in Figure 3D-E” and F). However, we could not rule out the possibility that the *Btk*^*SA11*^ spores altered EB behavior, pushing them toward an EEC fate. To verify this possibility, we carried out a ReDMM lineage tracing using the *Su(H)-Gal 4* driver that is specifically expressed in EBs (*Su(H)-ReDDM* flies). In this case, no newborn EECs were detectable upon ingestion of the *Btk*^*SA11*^ spores while many newborn ECs were present (Figure S3A-C), indicating that EBs are not the source of the increase in the number of EECs.

**Figure 3:**
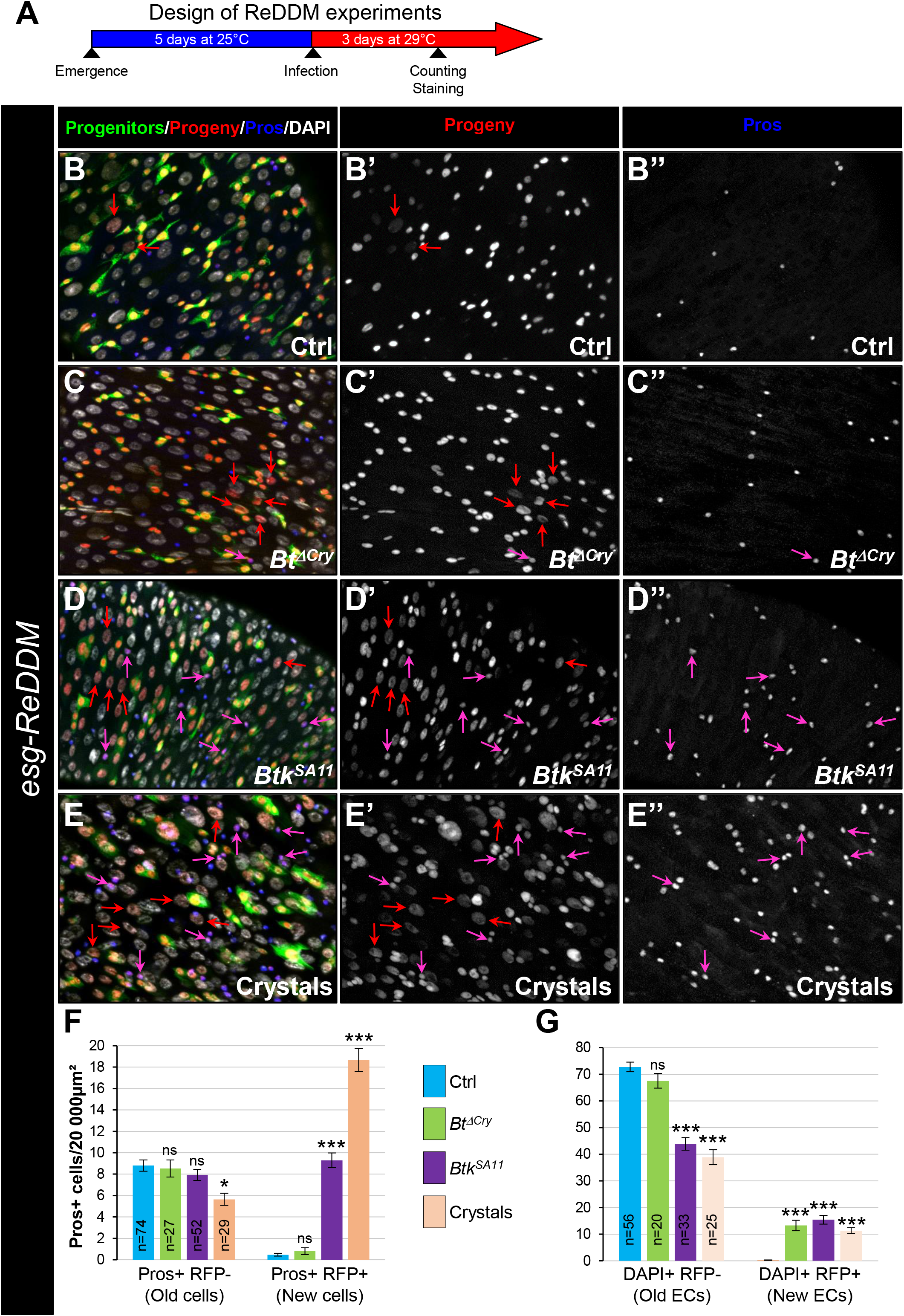
EEC excess arises from newborn EEPs after ingestion of *Btk*^*SA11*^ crystals. (**A**) Schema of the experimental design for the *esg-ReDDM* cell lineage used in this entire figure. (**B-E”**) R4 region of *esg-ReDDM* flies. Midguts were stained for Pros (blue) and DAPI which marks nuclei (white in B, C, D and E). (B-E’’) show the different cell types which either existed before the ingestion (green and red) or arise after the ingestion (red only) of water (B-B”, Ctrl), *BtkΔ*^*Cry*^ spores (C-C”) or *Btk*^*SA11*^ spores (D-D”) and Crystals (E-E’’). ISCs were GFP+ RFP+ DAPI+ with small nuclei; EBs were GFP+ RFP+ DAPI+ with bigger nuclei; EEPs were GFP+ RFP+ Pros+ DAPI+; new EECs were RFP+, Pros+ DAPI+; old EECs were Pros+ DAPI+; new ECs were RFP+ DAPI+ with polyploid big nuclei and old ECs were DAPI+ with very big nuclei. **(F)** Counting old EECs (Pros+ RFP-) and new EECs (Pros+ RFP+) in the conditions described in (B-E) **(G)** Counting old ECs (DAPI+) and new ECs (DAPI+ RFP+) in the conditions described in (B-E) n=number of samples; ns=not significant; * (P ≤ 0.05); *** (P ≤ 0.001).

Altogether, our data demonstrate that ingestion of *Btk*^*SA11*^ spores damages the intestinal epithelium, strongly stimulating symmetric ISC proliferation. However, some of the progenitors make the choice to commit to an EEP/EEC fate instead of an EB/EC fate. Consequently, there is a lack of new ECs to replace the dying ones and there is an excess of EECs. Interestingly, the crystals containing the Cry protoxins can recapitulate all the *Btk*^*SA11*^ spore effects. In contrast, the effect of *Btk*^*ΔCry*^ spores is less damaging for the gut epithelium. In this case, ISC proliferation is only weakly stimulated and the progenitors make the choice to commit to the EB/EC fate to replace the damaged ones.

### Crystals of Btk^SA11^ protoxins decrease ISC-Progenitor cell-cell adhesion

It is well established that the Notch (N) signaling pathway governs progenitor differentiation and cell lineage choice in the adult *Drosophila* midgut (Micchelli and Perrimon, 2006; Ohlstein and Spradling, 2006; Ohlstein and Spradling, 2007; Pasco et al., 2015). Indeed, the transmembrane ligand Delta (Dl) expressed in ISCs binds to its N receptor present on the surface of progenitors. This induces the cleavage of the intracellular domain of N and its relocation into the nucleus to activate its target genes (Perdigoto and Bardin, 2013). Upon N activation, progenitors differentiate into EBs and then into ECs while in the absence/weak activation of N signaling, progenitors commit to an EEP/EEC fate (Beehler-Evans and Micchelli, 2015; Guo and Ohlstein, 2015; Ohlstein and Spradling, 2007; Sallé et al., 2017). A prolonged and/or strong interaction between the ISC and its progenitor is necessary to reach the threshold of the N signaling activation sufficient to commit the progenitor to the EB/EC fate. A shorter and/or weaker interaction between the ISC and its progenitor weakly induces the N pathway, pushing the progenitor towards the EEC fate (Guisoni et al., 2017; Sallé et al., 2017). The adherens junctions between ISCs and progenitors formed by E-Cadherins and Connectins intervene to prolong the contact between Dl and N, favoring the EB/EC fate (Choi et al., 2011; Falo-Sanjuan and Bray, 2021; Maeda et al., 2008; Zhai et al., 2017). As the consensus receptors for Cry1A toxins in target organisms are members of the Cadherin family (Adang et al., 2014), we wondered whether the *Btk*^*SA11*^ could interfere with the function of the adherens junctions. We hypothesized that ISC-progenitor interaction could be reduced via interference of Cry toxins with Cadherins, modifying the progenitor cell fate and thus explaining the excess in EEC number seen after ingestion of *Btk*^*SA11*^ spores or purified crystals. To test this hypothesis, we labelled the intestines of *esg>GFP Drosophila* fed with *Btk*^*ΔCry*^ spores, *Btk*^*SA11*^ spores or purified crystals with the anti-Armadillo (Arm)/β-catenin antibody that strongly marks the adherens junctions. We observed an intense labeling at the level of the ISC-progenitor junctions in the control (Figure 4A-A’ and I) or following intoxication by *Btk*^*ΔCry*^ spores (Figure 4C-C’ and I). Strikingly, this labeling became less intense following ingestion of *Btk*^*SA11*^ spores or purified crystals (Figure 4E-E’, G-G’ and I). We confirmed these observations using a *Drosophila* line expressing the endogenous DE-Cadherin (DE-Cad) fused to a Tomato tag (Figure 4 B, D, F and H). Moreover, we observed that the decrease in Tomato::DE-Cad labelling between an ISC and its progenitor was correlated with the expression of EEP markers (blue stars in figure 4B and F and H). Our results suggested that crystals of Cry protoxins produced by *Btk*^*SA11*^ are responsible for the increase in the number of EEPs/EECs, this effect being associated with a decrease in intercellular adhesion between ISCs and progenitors.

**Figure 4:**
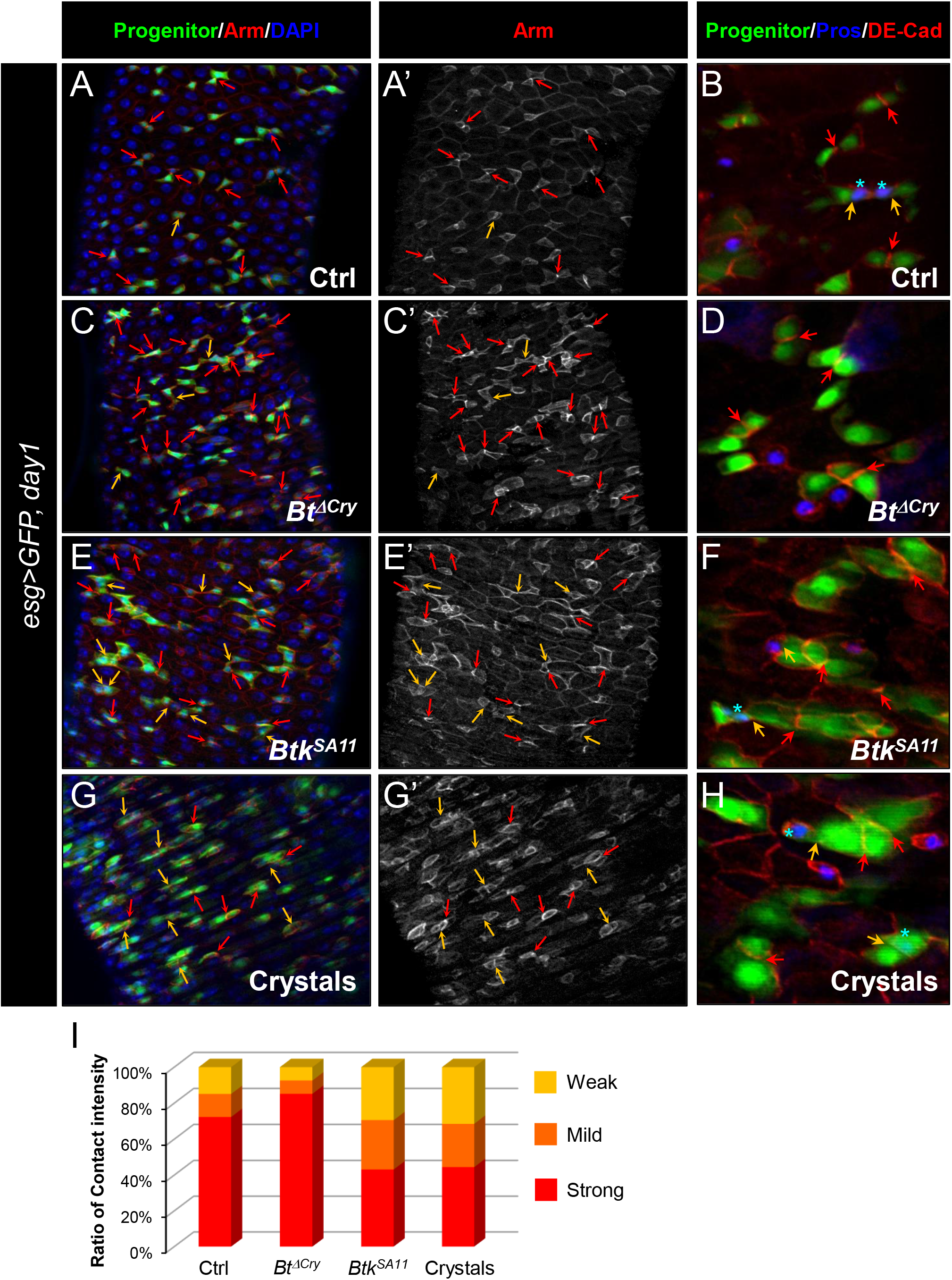
Btk bioinsecticide decreases ISC-Progenitor cell-cell adhesion. (**A-I**) *esg>UAS-GFP* or) *esg>UAS-GFP, Tomato::shg* (B, D, F and H) *Drosophila* midgut R4 region 24h PI of water (A-B Ctrl), *Btk*^*ΔCry*^ spores (C-D), *Btk*^*SA11*^ spores (E-F) or crystals (G-H). (**A, C, E**, and **G**) Midguts are labeled with anti-Armadillo (Arm) (red in A, C, E, and G) which strongly marks the adherent junctions between ISC and progenitors (green) and Dapi (blue). (**A’, C’, E’**, and **G’**) correspond to the single Arm channel. (**B, D, F** and **H**) Midguts are labelled for Pros (blue), DE-Cadherin (red) and ISCs and progenitors (green). Red arrows point to the high intensity of adherens junctions staining between ISC and progenitors. Yellow arrows point to the weak intensity of adherens junction staining. Note that the high intensity of adherens junction staining is associated with ISC/EB interaction while the weak intensity of adherens junction staining is associated with ISC/EEP interaction (blue stars mark EEPs in B, D, F and H) (**I**) Graph representing the percentage of the different categories of cell contact intensity between ISCs and progenitors in the experimental conditions shown in (A, C, E, and G). 15<n<18 intestines. Weak = Contact Intensity/Membrane Intensity<1.3; Mild = 1.3<Contact Intensity/Membrane Intensity<1.6; Strong = Contact Intensity/Membrane Intensity>1.6.

### Increasing adherens junction strength rescues crystal-dependent cell fate diversion

To confirm that the crystals of *Btk*^*SA11*^ interfered with progenitor fate by disturbing adherens junctions, we wondered whether increasing the strength of cell adhesion between ISCs and progenitors could rescue the right number of EEPs/EECs. Thus, we overexpressed the DE-Cadherin in these cells using the *esg-ReDDM* flies (Figure 5 and S4A-D’). We analyzed the identity of newborn cells 3 days after ingestion of water (control), *Btk*^*ΔCry*^ or *Btk*^*SA11*^ spores, or purified crystals. Interestingly, in flies overexpressing the DE-Cadherin in ISCs and progenitors fed with *Btk*^*SA11*^ spores or purified crystals, we no longer observed an increase in the number of EECs (blue arrows in Figure 5A-D, and Figure 5E). In agreement, the number of ISC-progenitor pairs with a strong interaction was increased (Figure 5F compared to Figure 4I). Furthermore, as expected, newborn ECs appeared (red arrows in Figure 5B-D’), strongly suggesting that increasing the cell adhesion between ISCs and progenitors rescued the progenitor fate disturbance generated by the *Btk*^*SA11*^ crystals. Surprisingly, overexpressing the Connectin, another cell adhesion molecule, which mediates hemophilic cell-cell adhesion (Zhai et al., 2017), in both ISCs and progenitors did not rescue the number of EECs following feeding with *Btk*^*SA11*^ spores or purified crystals (Figure S5). To confirm the disturbance of DE-Cadherin-dependent cell adhesion by purified crystals, we carried out an aggregation assay in *Drosophila* S2 cell culture. Indeed, S2 cells do not endogenously express the DE-Cadherin and display only a weak cell-cell adhesion phenotype (Toret et al., 2014). Transfection of S2 cells with a plasmid encoding a DE-Cadherin::GFP fusion resulted in large aggregate formation as early as 1h post-agitation (Ctrl in Figure 5G and S4E). Adding purified crystals to the cell culture medium strongly reduced the size of aggregates. Interestingly, purified Cry1Ab and Cry1Ac protoxins have the same effect although Cry1Ab needed a longer time to reduce the size of S2 cell aggregates (Figure 5G-G” and S4E). Because Cadherins serve as receptors for the Cry1A toxin family in target Lepidoptera (Adang et al., 2014), our data suggest that in non-target organisms such as *Drosophila melanogaster*, Cry1A toxins could interfere physically with the well-conserved E-Cadherin.

**Figure 5:**
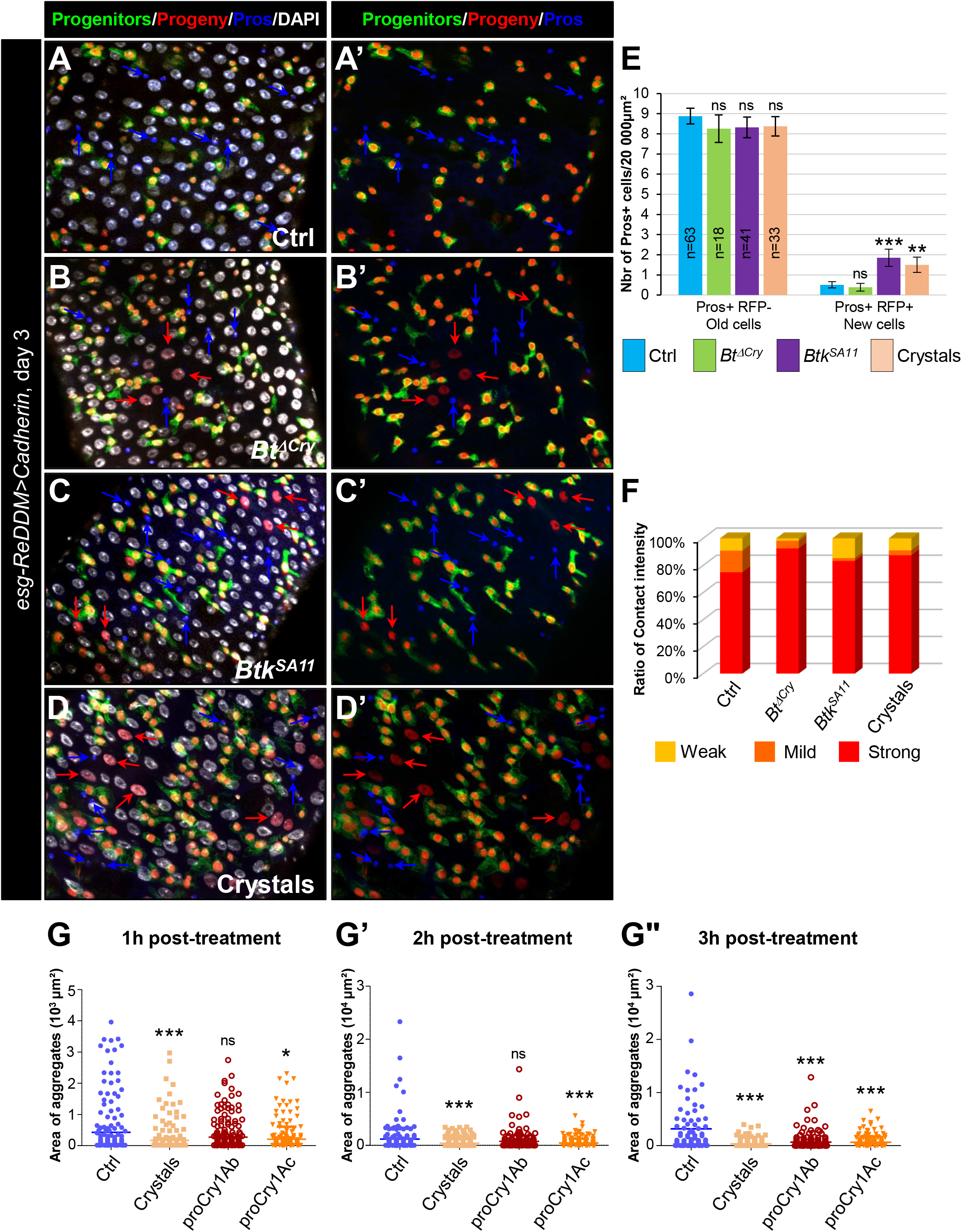
Increasing adherens junction strength rescues crystal-dependent cell fate diversion. (**A-F**) *esg-ReDDM>DE-Cad Drosophila* midgut R4 region. These flies specifically overexpress the DE-Cad in ISCs and progenitors. Flies were fed with water (A-A’, and blue in E, Ctrl), *Btk*^*ΔCry*^ spores (B, B’, and green in E), *Btk*^*SA11*^ spores (C, C’ and purple in E) or Crystal (D-D’ and pink in E) and observed 72h PI (see Figure 3A for the experimental design). In (A-D’) blue arrows point to old EECs and red arrows newborn ECs. (E) Number of old EECs (Pros+ RFP-) and new EECs (Pros+ RFP+) in the conditions described in (A-D). (**F**) Graph representing the percentage of the different categories of cell contact intensity between ISCs and progenitors in the experimental conditions shown in (Figure S4 A-D’). Weak = Contact Intensity/Membrane Intensity<1.3; Mild = 1.3<Contact Intensity/Membrane Intensity<1.6; Strong = Contact Intensity/Membrane Intensity>1.6. (**G-G”**) Cell aggregation assays on S2 cells expressing the DE-Cadherin::GFP. Cells placed under constant rotation were incubated with or without *Bt* crystals or purified Cry protoxins (Cry1Ab or Cry1Ac) for 1h (G), 2h (G’) or 3h (G”). Each scatter plot represents the area (µm2) of all objects (aggregates or individual cells) obtained from 3 independent experiments. Representative images of cell aggregates formed in aggregation assays are shown in supplemental data (Figure S4E). n=number of samples, ns (non-significant), * (P ≤ 0.05), ** (P ≤ 0.01), *** (P ≤ 0.001).

### Cry1A toxins mimic *Btk*^*SA11*^ spore effects

*Btk*^*SA11*^ produces 5 different Cry toxins (Cry1Aa, Cry1Ab, Cry1Ac, Cry2Aa, and Cry2Ab) (Caballero et al., 2020). We investigated whether the increase in EEP/EEC number was due to all toxins present in the crystals or to only one family of toxins (i.e., the Cry1A or Cry2A family). We made use of the *Btk*^*Cry1Ac*^ strain (referred to as 4D4 in http://www.bgsc.org/) which produces crystals composed only of the Cry1Ac protoxin. We fed *esg>GFP* flies either with spores of *Btk*^*Cry1Ac*^ or with purified Cry1Ac crystals. In both conditions, we observed a significant increase in the number of EEPs and EECs 3 days post-ingestion compared to controls (Figure 6A and S6A, C and D). To verify whether other toxins of the Cry1A family induced a similar rise in EEP/EEC number, we generated a *Btk*^*Cry1Ab*^ strain (see Material and Methods) producing only the Cry1Ab toxins (Figure S6I). Similar to the *Btk*^*Cry1Ac*^ spores, *Btk*^*Cry1Ab*^ spores induced an increase in EEP/EEC numbers (Figure 6B and S6B). Unfortunately, no *Btk* strain yielding only Cry2A-containing crystals was available and we were unsuccessful in generating one. To overcome this, we used heterologous expression of Cry toxins in *E coli*. We first checked whether Cry1Ac protoxin produced and purified from *Escherichia coli* (*E. coli*) was indeed able to induce an increase in EEP/EEC numbers. We also forced the activation of the Cry1Ac protoxin into an activated form *in vitro* (see Material and Methods). Interestingly, both the protoxin and the activated form of Cry1Ac were able to induce the expected phenotype, though the activated Cry1Ac form was more efficient (Figure 6C and Figure S6E and F). Conversely, both Cry2Aa protoxin and its activated form purified from *E coli* were unable to increase the number of EEP/EECs (Figure 6C and Figure S6G and H). Therefore, our data demonstrate that ingestion of Cry1A toxins was sufficient to induce a rise in the numbers of both EEPs and EECs.

**Figure 6:**
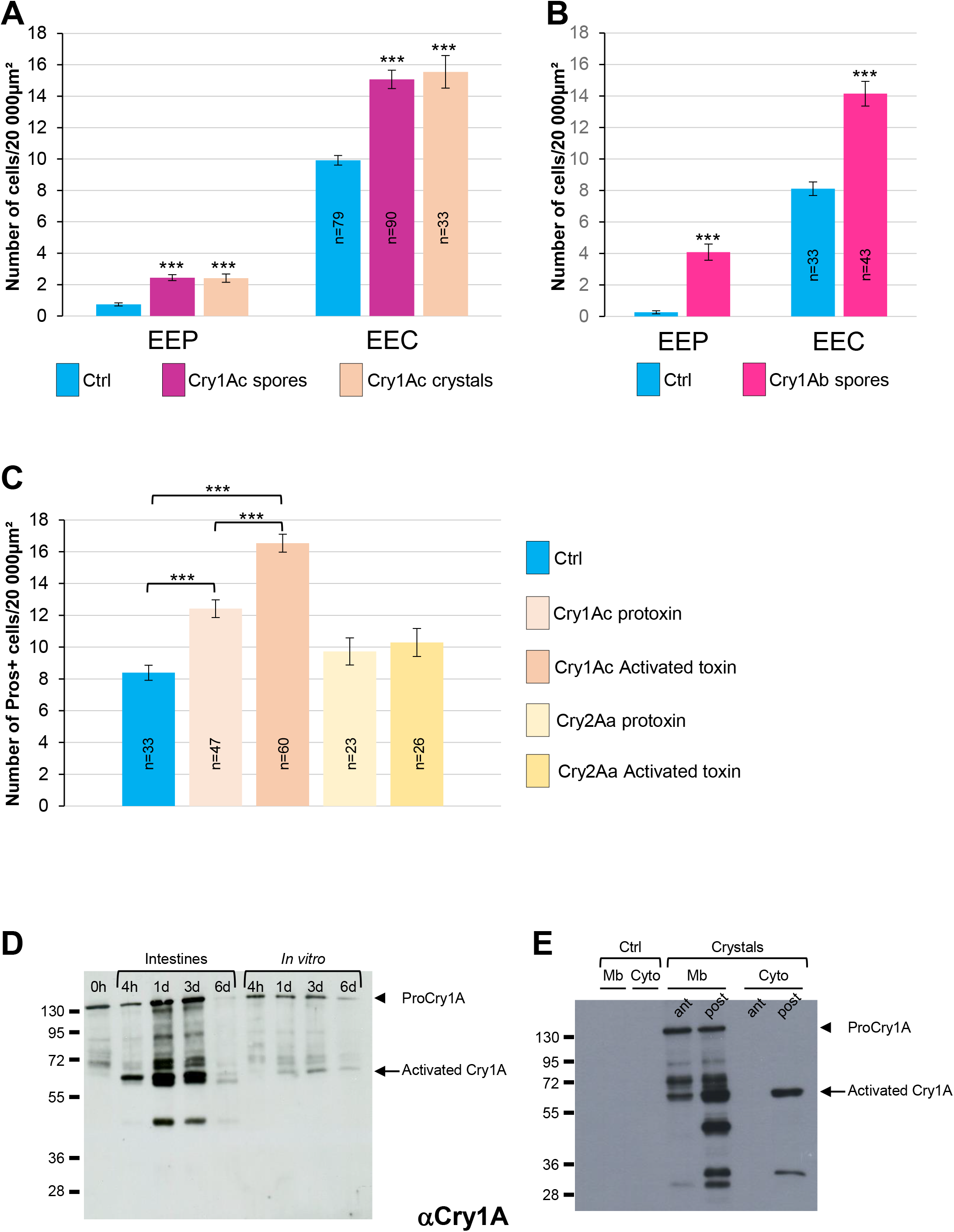
Cry1A toxins mimic *Btk* crystal effects. (**A-C**) *esg>GFP* flies fed with water (blue bars, Ctrl), *Btk*^*Cry1Ac*^ spores (fuchsia bars in A), Cry1Ac crystals (pink bars in A), *Btk*^*Cry1Ab*^ spores (fuchsia bars in B), Cry1Ac protoxins (light pink bar in C), Cry1Ac activated toxins (pink bars in C), Cry2Aa protoxins (light beige bar in C) and Cry2Aa activated toxins (beige bar in C). (**A** and **B**) Number of EEPs or EECs in the R4 region 72h PI **(C)** Number of Pros+ cells in the R4 region 72h PI (**D** and **E**) Western Blot from dissected intestines using a polyclonal Anti-Cry1A antibody detecting both the protoxins and the activated forms of Cry1A family of toxins. **(D)** (left lane) 0h corresponds to *Btk*^*SA11*^ spores extemporaneously resuspended in water. (Right part of the blot) *Btk*^*SA11*^ spores from incubated *ex vivo* (control) in water at 25°C for the period of the experiment. We mainly detect the protoxin forms of Cry1A at 130kDa (arrowhead). (Left part of the blot) Proteins extract from midgut of flies fed by the same *Btk*^*SA11*^ preparation (T 0h) at 4h and 1, 3 and 6 days PI. The 130kDa protoxins are still visible. The 67kDa activated form appears as early as 4h (arrow). 6 days PI no more toxins are detected in the midgut. **(E)** Flies fed 2 days with water (Ctrl, left part) or with purified crystals (right part). Protoxins (130kDa) are present in the membrane fraction (Mb) in both the anterior (ant) and the posterior (post) midgut. The 67kDa activated forms are present in the membrane fraction of both the anterior and posterior midgut and in the cytoplasmic fraction (Cyto) of the posterior midgut. n=number of samples. *** (P ≤ 0.001).

### Cry1A Protoxins from *Btk* crystals are activated in the *Drosophila* midgut

Our data above showed that purified activated Cry1Ac toxin was more efficient for inducing an EEP/EEC excess than the purified Cry1Ac protoxin. Interestingly, the magnitude of EEP/EEC excess was similar using either Cry1Ac crystals or purified activated Cry1Ac toxin (compare Figure 6A and 6C), suggesting that Cry1Ac protoxins contained in the crystals were activated in the *Drosophila* intestine. However, the admitted model proposes that protoxins can be activated *in vivo* only in the intestine of the susceptible lepidopteran owing to the presence of appropriate digestive proteases specifically functioning at the basic pH and reducing conditions encountered in the larval midgut of lepidopteran (Pardo-Lopez et al., 2012; Soberon et al., 2009; Vachon et al., 2012). We therefore wondered whether the effects we observed *in vivo* were due to the crystal on its own (i.e., protoxins) or to the activated Cry toxins after processing in the *Drosophila* midgut. We first monitored by Western Blot the processing of the Cry1A toxin family in the fly midgut fed with *Btk*^*SA11*^ spores. We used an anti-Cry1A antibody raised against the activated form of the toxin, which therefore recognizes both forms (Babin et al., 2020). As a control, we incubated *Btk*^*SA11*^ spores *in vitro* in water, at 25°C for 4h, 1d, 3d and 6d. Under these conditions, the protoxin form of Cry1A at 130kD was predominant and stable for at least 3 days before fading (control in Figure 6D, right part of the blot). This observation is in agreement with the fact that the half-life of Cry1A crystals has been estimated at about 1 week in soil or under laboratory conditions at 25°C (Hung et al., 2016). We used the same initial *Btk*^*SA11*^ spore preparation (0h) to feed the flies. We further dissected intestines and extracted total proteins at different times post-feeding. Interestingly, as early as 4h, we observed the 67kD activated form of the Cry1A toxins (Figure 6D left part). Noteworthy, the 130kD protoxin forms were still present, as the flies kept ingesting spores and crystals throughout the experiments. Six days after ingestion, almost no more protoxins or toxins were detectable due to the instability of the crystals (Figure 6D). Thus, our data show that the crystals can be processed in the midgut of adult *Drosophila* to give rise to active forms of the Cry1A toxins.

As mentioned previously, the increase in EEPs/EECs numbers was prominent in the posterior midgut (Figure S2). We therefore wondered whether Cry1A crystals were differentially processed along the midgut. We fed flies with purified *Btk*^*SA11*^ crystals and 24h later, we dissected and crushed the intestines by separating the anterior midgut from the posterior part. Furthermore, we performed subcellular fractioning by separating the soluble fraction (considered as the cytosol-enriched fraction) and the insoluble fraction containing membranes. Interestingly, we observed that protoxins were already activated in the anterior part of the midgut (the 67kD form) but were only associated with membranes (Figure 6E). In the posterior midgut, we detected a stronger quantity of the 67kD-activated form both at the membrane and in the cytosol, suggesting an internalization by epithelial cells (Figure 6E). In this part of the midgut, smaller forms appeared mainly associated with the membrane, probably resulting from degradation processes. Notably, we did not detect the 130kD protoxins in the cytoplasmic fraction of both anterior and posterior midguts, suggesting that the protoxins remained associated with membranes. Moreover, total processing was never achieved since some protoxins still remained detectable. Altogether, our data show that crystals containing Cry1A protoxins are processed all along the adult *Drosophila* midgut to generate activated Cry1A toxins.

## Discussion

Our results show that the Cry1A toxin family of *Btk* disrupts the gut cellular homeostasis of the non-susceptible organism *Drosophila melanogaster*. Cry1A induces EC death coupled to an increase in ISC proliferation to replace the damaged ECs. Importantly, Cry1A toxins also altered intestinal cell composition by weakening DE-Cadherin-dependant cell-cell adhesion which is normally highly enriched in adherens junctions linking ISCs to their immediate progenitors (Choi et al., 2011; Maeda et al., 2008; Ohlstein and Spradling, 2006; Zhai et al., 2017). As a consequence, progenitors are pushed toward the EEC path of differentiation instead of the EC, owing to reduced activation of N signaling in progenitors (Figure 7) (Guo and Ohlstein, 2015; Maeda et al., 2008; Ohlstein and Spradling, 2006; Ohlstein and Spradling, 2007; Pasco et al., 2015; Zhai et al., 2017). Our data confirmed that the duration and/or the strength of the cell-cell contact between the ISC and its progenitor daughter cell is important to drive the progeny toward an appropriate fate. Indeed, it has been shown both in *Drosophila* and in mammalian cell culture that adherens junctions are crucial to reinforce the contact between neighboring cells to allow the activation of N signaling (Falo-Sanjuan and Bray, 2021; Shaya et al., 2017; Zhai et al., 2017), N being necessary for the EB-EC cell fate choice.

**Figure 7:**
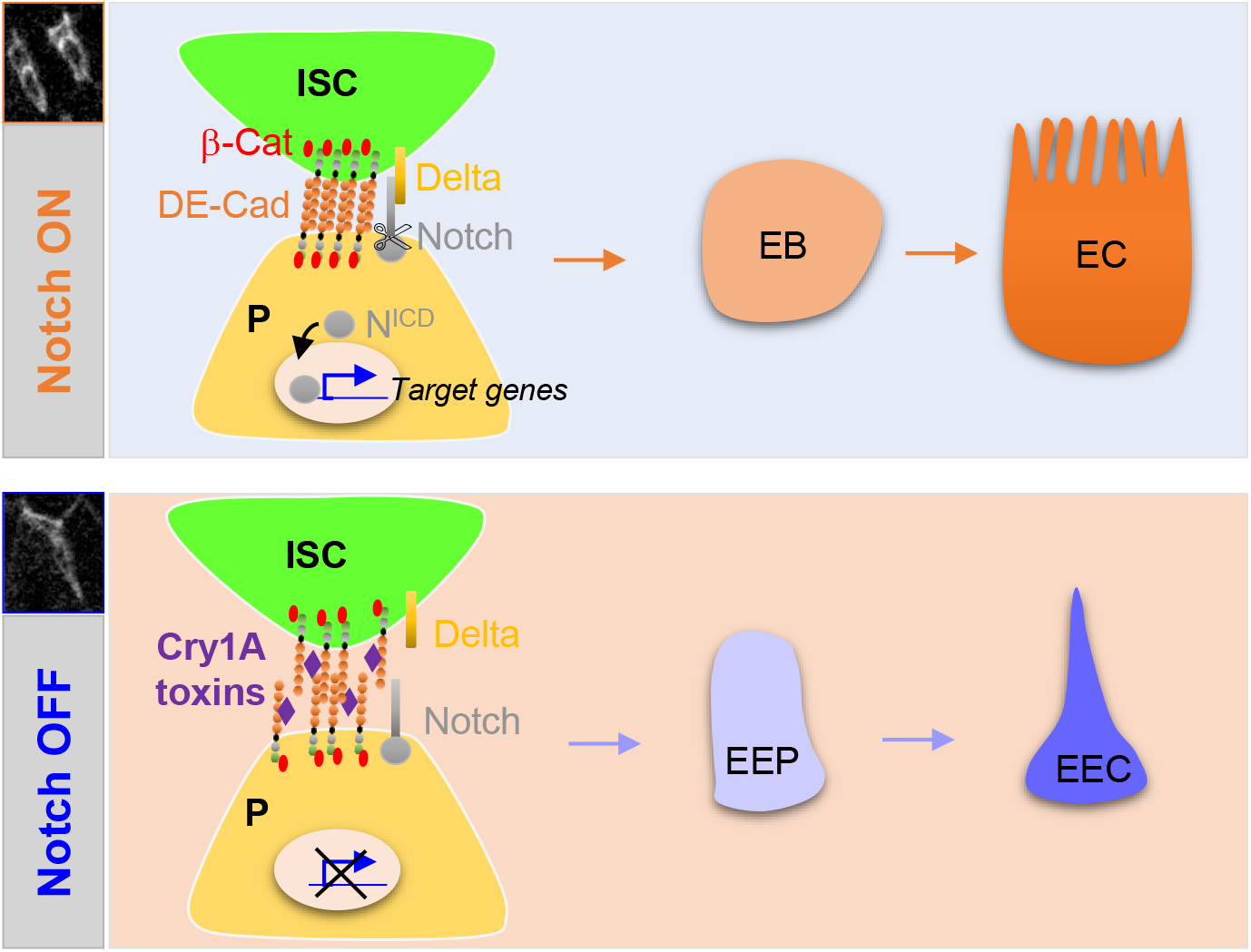
Cry1A toxins interfere with progenitor fate behavior. **Notch ON:** in *Drosophila*, 90% of ISC daughter cells commit to the EB/EC fate owing to the strong activation of the Notch signaling pathway in the EBs. The adherens junction DE-Cadherin (DE-Cad)-dependent are required to permit the interaction between the Delta ligand in ISC and the Notch receptor in EB. **Notch OFF:** Ingestion of Cry1A toxins impedes the DE-Cad homophilic interaction between the ISCs and their progenitor daughter cells, reducing the activation of Notch signaling in progenitors. Consequently, progenitors adopt an EEP/EEC fate.

Only the Cry1A family of toxins induces an increased number of EECs while Cry2A toxins do not display any phenotype. Previous studies showed that Cry1A and Cry2A bind to different receptors in the intestinal epithelium of susceptible lepidopteran larvae, and proteins of the Cadherin family appear essential for Cry1A, but not Cry2A, binding (Adang et al., 2014; Gao et al., 2019; Hernández-Rodríguez et al., 2013; Li et al., 2020; Liu et al., 2018a). *Drosophila* possesses 17 genes encoding for members of the Cadherin superfamily (Hill et al., 2001). Cad88C is the most similar to Bt-R Cadherin of susceptible lepidoptera, but shares only 17% of identity, (Stevens et al., 2017) and is poorly expressed in the *Drosophila* midgut (see http://flygutseq.buchonlab.com/ and https://flygut.epfl.ch/). However, Cry1A effects on *Drosophila* progenitor cell fate appear to depend specifically on the DE-Cadherin present in the adherens junctions, since overexpression of DE-Cadherin in progenitor cells (ISCs, EBs and EEPs) can overcome Cry1A-induced impacts while another component of the adherens junctions (Connectin) cannot. Interestingly, the percentage of identity between the Cry1A binding regions (CBRs) shared by the orthologs of Cadherin-type receptors in different susceptible lepidopteran species ranges from 21% to 66% (Li et al., 2021; Shao et al., 2018). Hence, the presence of a well-conserved primary consensus sequence within the CBRs cannot explain the specificity of binding. In agreement, it has been recently shown in susceptible Lepidoptera that only two dipeptides within the CBRs are essential for high-affinity binding of Cry1A to its Cadherin receptor (Liu et al., 2018a). These two dipeptides are not conserved in the CBRs of the Bt-R orthologs in different lepidopteran species targeted by Cry1A toxins (Li et al., 2021). These data suggest that binding of Cry1A to the receptor relies more on a conserved conformation of the CBRs than on a conserved primary sequence. In addition, alignment of the *Helicoverpa armigera* (a Cry1A susceptible lepidopteran) CBR sequence (Gao et al., 2019) and the DE-Cadherin sequence between amino acids 164 and 298 shows 24% identity and 47% similarity (using BlastP, https://blast.ncbi.nlm.nih.gov/). Altogether, these results support a possible binding of Cry1A toxins to *Drosophila* intestinal DE-cadherins, although this could occur with a low affinity.

Susceptibility of lepidopteran larvae to CryA1 toxins relies on the presence of a secondary receptor such as Alkaline phosphatases, Aminopeptidases N, or ABC transporters (Adang et al., 2014; Gao et al., 2019; Li et al., 2020; Liu et al., 2018b). None of the orthologs of these receptors has been shown to be strongly expressed in the *Drosophila* midgut (Li et al., 2021; Stevens et al., 2017), which could explain why binding of Cry1A to DE-cadherins does not lead to the death of adult *Drosophila*. Interestingly, in *Drosophila* larvae, heterologous expression of ABCC2 from *Bombix mori* or *Plutella xylostella*, or Aminopeptidases N from *Manduca sexta* was sufficient to induce Cry1A-dependent death (Gill and Ellar, 2002; Obata et al., 2015). Conversely, heterologous expression of the *Bombix mori* Cadherin Bt-R receptor in *Drosophila* is not sufficient to induce death upon exposure to Cry1A but strongly enhances death when co-expressed with BmABCC2 (Obata et al., 2015). Together, these data suggest that, in *Drosophila*, in the absence of a Cadherin receptor displaying a high affinity to Cry1A toxin, endogenous Cadherins with reduced affinity toward Cry1A could step in to enhance the death potential of Cry1A toxins in the context of heterologous expression of a secondary receptor from susceptible Lepidoptera. In agreement, it has been previously demonstrated in *Drosophila* larvae that increasing the dose of ingested *Btk* spores and crystals could ultimately lead to death (Babin et al., 2020; Cossentine et al., 2016), arguing that the receptors (i.e., the Cadherins and the secondary receptors) present in the *Drosophila* midgut have lower affinities for Cry1A toxins.

Cry toxin activities in susceptible organisms rely not only on the presence of specific host receptors in the midgut, but also on the extreme midgut pH and reducing environment allowing crystal solubilization, as well as the enzymatic capacity of digestive proteins, both of which are involved in the conversion of protoxins into active toxins (Fiuza et al., 2017; Mendoza-Almanza et al., 2020; Pardo-López et al., 2013; Shao et al., 2013). Nevertheless, our data along with another recent study (Stevens et al., 2017) suggest that crystals can be solubilized and then protoxins activated in the *Drosophila* midgut. Therefore, more investigations will be necessary to understand how crystals are processed and protoxins activated in the intestine of non-susceptible organisms. What makes the difference between Cry-susceptible and non-susceptible organisms is likely the affinity of Cry toxins for midgut host receptors. The higher the affinity of the toxins, the greater the cellular damage/death and the greater the risk of death. In addition, the capacity of regeneration of the midgut epithelium also plays an important role to overcome *Bt* pathogenicity (Castagnola and Jurat-Fuentes, 2016). If the regeneration of the intestinal epithelium is more efficient than the destructive capacity of Cry toxins, the host will survive. However, whatever the host is (susceptible or non-susceptible), *Bt* uses a three-pronged strategy to improve its degree of virulence. First, it tries to disrupt the epithelial barrier function by damaging or killing ECs. Second, it diverts the behavior of the progenitor cells toward the wrong fate, thus limiting the amount of ECs produced that are necessary to replace the damaged ones and to maintain the midgut integrity. Third, killing ECs also reduces the capacity of these cells to produce reactive oxygen species and antimicrobial peptides known to be involved in antimicrobial defenses (Allaire et al., 2018; Capo et al., 2016; Kim and Lee, 2014).

The *Bt* subspecies *kurstaki* and *aizawai* are widely used as microbial pesticide to fight lepidopteran pests worldwide (Casida and Bryant, 2017), both subspecies partially producing the same Cry1A toxins (Caballero et al., 2020). Likewise, 180 millions of hectares of genetically modified crops express Cry1A toxins. Consequently, Cry1A toxins are present in the food, the feed and in the environment, implying that many organisms might be affected. As the mechanisms of intestinal progenitor fate choice are conserved in the animal kingdom (Guo et al., 2021; Joly and Rousset, 2020; Zwick et al., 2019), it would be interesting to investigate whether Cry1A toxins can also promote an increased number of EECs in other organisms (vertebrates and invertebrates). EECs, through the production of neuropeptides and hormones, are involved in the regulation of many physiological functions such as feeding behavior, metabolism and immune response (Guo et al., 2021; Nässel and Zandawala, 2019; Watnick and Jugder, 2019). Consequences of this increase in EEC number could be, for example, metabolic dysfunctions or inflammatory pathologies. More studies are needed to understand the physiological impacts of this change in intestinal cellular composition on organismal health.

## Materials and Methods

### Bacterial Strains

*Btk*^*ΔCry*^ (identified under the code 4D22, (Gonzalez et al., 1982)), *Btk*^*Cry1Ac*^ (identified under the code 4D4), and the *E. Coli* strains producing Cry1Ac (identified under the code ECE53), Cry1Ab (identified under the code ECE54) and Cry2Aa (identified under the code ECE126) were obtained from the Bacillus Genetic Stock Center (www.bgsc.org). The strain *Btk* SA-11 (*Btk*^*SA11*^) was isolated from the commercial product Delfin.

### Generation of Btk^Cry1Ab^

The mutant *Btk*^*Cry1Ab*^ producing only Cry1Ab as crystal toxin was obtained from the WT strain *Btk*^*SA11*^, by a procedure of plasmid curing, as follows. After isolation on TSA-YE agar (Biomérieux, 18h culture at 30°C), the strain *Btk*^*SA11*^ was sub-cultured successively 3 times in 10ml of brain heart Infusion (BHI, Oxoid) broth at 42°C with agitation, for 64, 48 and 36h respectively. The first BHI culture was inoculated from an isolated colony, and the subsequent cultures were inoculated with 100 µl of the previous ones. Clones from the last culture were isolated on TSA-YE agar after serial dilution, then subcultured on the sporulating medium hydrolysate of casein tryptone (HCT) + 0.3% Glc, in order to verify the absence of crystal production. Initially seeking to select clones totally free of cry genes, we fortuitously selected partial mutants, including this strain for which the single presence of *cry1Ab* as *cry* toxin gene, could be confirmed after WGS analysis, as follows. The genomic DNA of *Btk*^*Cry1Ab*^ and *Btk Btk*^*SA11*^ was extracted using the KingFisher cell and Tissue DNA kit (ThermoFisher) and sequenced using Illumina technology at the Institut du Cerveau et de la Moelle Epinière (ICM) platform, as previously described (Bonis et al., 2021), (NCBI accession numbers SAMN23436140 and SAMN23455549, respectively). The absence of *cry* genes in *Btk*^*Cry1Ab*^, with the exception of *cry1Ab*, was confirmed from raw reads using KMA (Clausen et al., 2018).

### Spore production

From isolated colonies on LB agar Petri dish, 4 × 5mL of *Bt* pre-culture was carried out. The pre-culture was used for sowing 4 × 500mL of PGSM medium (0.75% casamino acids, 0.34% KH_2_PO_4_, 0.435% K_2_HPO_4_, 0.75% glucose, 1.25mM CaCl_2_, 0.123% MgSO_4_, 0.002% MnSO_4_, 0.014% ZnSO_4_, 0.02% FeSO_4_) and allowed to grow and sporulate in an incubator shaker at 30°C, 180 rpm for 2 weeks. In order to eliminate vegetative cells, culture was heated 1 hour at 70°C and then centrifuged 15 min at 7500g. The pellet was resuspended with a Dounce homogenizer in 1L of 150 mM NaCl and placed for 30 min on roller agitation at room temperature. After centrifugation (15 min, 7500g), the pellet was washed twice with sterile water. The final pellet was resuspended in 30mL of sterile water, dispatched in 1mL weighed tubes and lyophilized for 24-48h. The spore mass was determined by the difference between the full and the empty tubes’ weights.

### Spore titer

The lyophilized spores were resuspended in sterile water to obtain a concentration of 50 mg/ml. This solution was diluted serially (100µl in 900 µl of sterile water) to obtain 10^−1^ to 10^−9^ dilutions. 100 µl of dilutions 10^−5^ to 10^−9^ were plated on LB agar and incubated at 30°C overnight. The number of colonies was counted for each dilution and reported to the mass of spores plated. The experience was renewed three times. The mean of these ratios allows us to determine the titer of spores in CFU (colony forming unit)/g.

### Crystal purification

2×1g of *Btk*^*SA11*^ lyophilized spores were resuspended in 2 × 30 mL of sterile water and placed on roller agitation at 4°C for 5h. Then, the solution was sonicated for 4 cycles of 15 sec/15 sec with a frequency of 50% (Fisherbrand™ Model 505 Sonic Dismembrator). 6 × 10 mL were deposited on a discontinuous sucrose gradient (67%/72%/79%) and centrifuged overnight in SW28 swinging buckets at 100 000g at 4°C (ultracentrifuge Thermo-Sorval WX Ultra 80). The crystals were collected at the 72%/79% and 72% /67% interfaces with a micropipette and dispatched by 10 mL in centrifuge tubes (Beckman Avanti JE, rotor JA 25.50). 25 mL of sterile water were added in each tube, vortexed and centrifuged at 4°C at 16 000g for 15 min. Each pellet was resuspended with 20 mL of sterile water and centrifuged again in the same conditions. Each final pellet was resuspended in 2 mL of sterile water, aliquoted by 1ml in weighed microtubes and lyophilized 24h-48h. The crystal mass was determined by the difference between the full and the empty tubes’ weights.

### Cry protoxin production

2 × 15 mL of LB ampicillin (50mg/mL) were inoculated with two colonies of *E. Coli* expressing the desired Cry toxin, and allowed to grow at 37°C, 220 rpm overnight. 4 × 5 mL of the overnight preculture were added to 4 × 500 mL of LB ampicillin (50mg/mL) and allowed to grow at 37°C and 200rpm until DO_600_=0,6-0,7. Cry protein expression was induced by adding 500 µL of 1M IPTG in each culture. The cultures were left at 37°C and 200rpm overnight and then centrifuged at 6500g, 15 min at 22°C. Pellets were pooled in two batches and resuspended 100 mL of cold WASH buffer (20mM Tris, 10mM EDTA, 1% Triton X-100, pH 7,5) and incubated 5 min at 4°C. 500 µL of lysozyme (20mg/mL) were added in each solution and incubated 15 min at 30°C and then 5 min at 4°C. The solutions were sonicated for 6 cycles of 15 sec/15 sec’ at 40% (Fisherbrand™ Model 505 Sonic Dismembrator) and centrifuged 10 min at 10 000g, at 4°C. The pellets were washed twice with 100ml of WASH buffer and centrifuged in the same conditions. The last pellets were weighed, resuspended in CAPS buffer (50mM CAPS, 0,3% lauroyl-sarcosine, 1mM DTT pH11) to obtain a final concentration of 100mg/mL, placed for 30 min under roller agitation at room temperature and centrifuged 10 min at 10 000g, 4°C. The supernatant was dialyzed twice against 50 volume of PBS1x, 0,1mM DTT and twice against 50 volume of PBS1x at 4°C and centrifuged 10 min at 20 000g at 4°C. The supernatant was conserved at -20°C until purification or digestion (activation) by trypsin.

### Cry toxin activation

Half of the produced supernatant (see above) was dosed by the Bradford method (Protein Assay Dye Reagent Concentrate, Biorad #500-0006) and digested with 1% trypsin (weight/weight) (Trypsin from bovine pancreas, Sigma #T1005) at 37°C for 72h. The Cry toxins were then purified by FPLC. NB: After 72h, the trypsin is fully degraded.

### Cry toxin purification

The Cry toxins produced from *E.Coli* (activated or not) were purified by FPLC (Äkta, UPC900/P920/INV907/FRAC950) on a 1 mL benzamidine column (HiTrap™ Benzamidine FF (high sub), GE Healthcare #17-5143-01) with PBS1X as charge buffer, PBS1x, 1M NaCl as buffer for non-specific link and 100mM glycin pH3 as elution buffer.

### Fly Stocks and Genetics

The following stocks are listed at the Bloomington *Drosophila* Stock Center (https://bdsc.indiana.edu/): WT canton S (#64349). *w; Sco/CyO; tub-GAL80*^*ts*^*/TM6b* (#7018). *w; tub-GAL80*^*ts*^; *TM2/TM6b* (#7019). *w; esg-GAL4*^*NP5130*^ (#67054). *w; UAS-GFP/TM3 Sb* (#5430). *w; UAS-shg-R* (DE-Cadherin)(#58494); *y w, shg::Tomato* (#58789).

#### Other stocks

*w;* ; *Dl-GAL4/TM6b* (Zeng et al., 2010). *w; tub-GAL80*^*ts*^; *Dl-GAL4 UAS-GFP/TM6b* (this study). *w; esg-GAL4*^*NP5130*^ *UAS-GFP* (Shaw et al., 2010). *w; esg-GAL4*^*NP5130*^ *UAS-GFP; tubGAL80*^*ts*^ (Apidianakis et al., 2009). *w; Su(H)GBE-GAL4, UAS-CD8::GFP* (Zeng et al., 2010). *w; Su(H)GBE-GAL4/SM6β; tub-GAL80*^*ts*^ *UAS-GFP/TM6b* (this study). *w; myo1A-GAL4* (Shaw et al., 2010). *w; myo1A-GAL4 UAS-GFP/CyO* (Apidianakis et al., 2009). *w; UAS-GFP::CD8; UAS-H2B::RFP/TM2* (Antonello et al., 2015). *w; UAS-CD8::GFP; UAS-H2B::RFP, tub-GAL80*^*ts*^*/TM2* (Antonello et al., 2015). *w; esg-GAL4, UAS-CD8::GFP; UAS-H2B::RFP, tub-GAL80*^*ts*^*/TM6b* (Antonello et al., 2015). *w; UAS-CD8::GFP; Dl-GAL4, UAS-H2B::RFP/TM6b* (this study). *w;* ; *UAS-GC3Ai*^*G7S*^ *(UAS-Casp::GFP)* (Schott et al., 2017). *w; UAS-connectin* (Zhai et al., 2017).

### Cell lineage

#### *Dl-ReDDM* experiments

*w; tub-GAL80*^*ts*^; *Dl-GAL4 UAS-GFP/TM6b* females were crossed to *w; UAS-GFP::CD8; UAS-H2B::RFP/TM6* males at 18°C. Progeny were kept at 18°C until emergence. Flies were transferred at 25°C for 5 days before infection, and then transferred at 29°C for 2 days (Figure S1B).

#### *esg-ReDDM* experiments

*w; esg-GAL4 UAS-GFP; UAS-H2B::RFP, tub-GAL80*^*ts*^*/TM6b* females were crossed to WT males at 18°C. Progeny were kept at 18°C until emergence. Flies were transferred at 25°C for 5 days before infection, and then transferred at 29°C for 3 days (Figure3).

#### *Su(H)-ReDDM* experiments

*w; Su(H)-GAL4/SM6β; tub-GAL80*^*ts*^ *UAS-GFP/TM6b* female*s* were crossed to *w; UAS-GFP::CD8; UAS-H2B::RFP/TM6* males at 18°C. Progeny were kept at 18°C until emergence. Flies were transferred at 25°C for 5 days before infection, and then transferred at 29°C for 3 days (Figure S3).

#### DE-Cadherin and Connectin overexpression

*w; esg-Gal4 UAS-GFP; UAS-H2B::RFP, tub-GAL80*^*ts*^*/TM6b* females were crossed to *UAS-DE-Cadherin* males. Progeny were kept at 18°C until emergence. Flies were transferred at 25°C for 5 days before infection, and then transferred at 29°C for 3 days (Figure 5 and S5).

### Drosophila rearing and oral infection

*Drosophila* were reared on standard medium (0.8% Agar, 2.5% sugar, 8% corn flour, 2% yeast) at 25 ° C with a 12h light/12h dark cycle. For oral infection, after 2 h of starvation to synchronize the food intake, ten 5 to 6 day-old non-virgin females were transferred onto a fly medium vial covered with a filter disk soaked with water (control) or a suspension of spores, (corresponding to 10^6^ CFU of spores per 4 cm² and per individual female (Loudhaief et al., 2017; Nawrot-Esposito et al., 2020)), crystals, protoxin or activated toxins. The quantity of crystals, protoxin or activated toxins deposited on the filter disc corresponded to 30% of the spore weight, *Btk* crystals representing between 25% and 30% of the total weight of the 1:1 spore/crystal mix (Agaisse and Lereclus; 1995; Monroe, (1961); Murty et al., 1994). Flies were kept feeding on the contaminated media until dissection in all the experiments.

### Dissection, immunostaining and microscopy

Dissection, fixation and immunostaining were performed as described by (Micchelli, 2014). Dilutions of the various antibodies were: mouse anti-Armadillo N27A1 at 1:50 (DSHB), mouse anti-Connectin-C1-427 at 1/200 (DSHB), mouse anti-Prospero MR1A at 1:200 (DSHB), rabbit anti-PH3 at 1:1000 (Millipore, 06-570), Rabbit anti-Cleaved Caspase-3 at 1/600 (Cell Signalling Asp175 #9661), Goat anti-mouse AlexaFluor-647 at 1/500 (Molecular Probes Cat# A-21235), Goat anti mouse AlexaFluor-546 at 1/500 (Molecular Probes Cat# A-11003), Goat anti-rabbit AlexaFluor-647 at 1/500 (Thermo Fisher Scientific Cat# A32733), Goat anti-rabbit AlexaFluor-546 at 1/500 (Thermo Fisher Scientific Cat# A-11010). For microscopy, guts were mounted in Fluoroshield DAPI medium (Sigma, # F6057) and immediately observed on a Zeiss Axioplan Z1 with Apotome 2 microscope. Images were analyzed using ZEN (Zeiss), ImageJ and Photoshop software. Image acquisition was performed at the microscopy platform of the Sophia Agrobiotech Institute (INRAE1355-UCA-CNRS7254 – Sophia Antipolis).

### DNA constructs

The full-length expression construct of DE-cadherin Full Length fused to GFP (DEFL) was introduced into pUAST as previously described (Oda and Tsukita, 1999).

### Cell aggregation assay

*Drosophila* Schneider 2 (S2) cells were cultured in Schneider’s medium supplemented with 10% heat-inactivated fetal bovine serum (FBS) at 25°C in a non-humidified ambient air-regulated incubator. For S2 aggregation assay, 2.2 10^6^ cells were plated in 25 cm^2^ flask for each condition. After 6 h, transient transfection was performed by mixing transfection reagent (TransIT®-2020; Mirus Bio) with a reagent-to-DNA ratio of 3:1. A total of 3 µg plasmid DNA per T25 was used, corresponding to a 5:1 mixture of pUAST-DEFL and pWA-GAL4. Approximately 46h after transfection, the cells were collected into 15 mL tubes and centrifuged for 5 minutes at 400 g. The pellet was resuspended in 2 ml of fresh medium supplemented in CaCl_2_ to obtain a final concentration of 7.4 mM and separated into single cells by repeated pipetting. 500 µL of this cell suspension were added to a well of a 24-well microplate. To allow cell-cell adhesion (aggregation), the microplate was placed under constant agitation on a rotary platform at 150 rpm at 25°C for the indicated time (1h, 2h and 3h) with or without protoxins (at a final concentration 35µg/ml). Cell aggregates formed in the wells were observed using an inverted Fluorescent microscope (Nikon, Eclipse TE2000-U). Images for the florescence of GFP was acquired using a CCD camera (ORCA ER, Hamamatsu Photonics). The same parameter settings were used to acquire images (objectives, gain, exposure time …). The area of fluorescent aggregates and individual cells were measured using Fiji software (Schindelin et al., 2012). The average area of a S2 cell in our condition is about 15.5 µm^2^. Hence, quantification of the aggregates area was performed excluding all objects smaller than 15 µm^2^. The mean area of aggregates were calculated after background subtraction. Three independent experiments were performed for each condition. Values in µm^2^ were represented in GraphPad Prism 5 software as scatter-plot view. Statistical analysis was conducted using GraphPad Prism 5 software. The significance of the difference between CTR and exposed conditions was assessed using one-way ANOVA and Tukey’s post hoc tests. Statistical parameters for each experiment can be found within the corresponding figure legends.

### Western blot

Figure 6D: Twenty *Drosophila* (5 to 6 day-old non-virgin females) were orally infected with spores and reared at 25°C with a 12h/12h day/night cycle for the indicated time. At the same time, 10^7^ CFU of spores in 50µl of sterile water were incubated in the same conditions. After 4, 24, 72 or 144h, *Drosophila* midguts were dissected in PBS1x with anti-proteases (cOmplete Tablets EDTA free EASY Pack, Roche #04693132001). Then midguts were transferred on ice into a 2 ml microtube containing 200 µL of PBS1x with anti-proteases and crushed one minute at 30Hz with a Tissue Lyser (Qiagen, Tissue Lyser LT).

Figure 6E: Twenty *Drosophila* (5 to 6 day-old non-virgin females) were orally infected with water or purified crystals and reared at 25°C with a 12h/12h day/night cycle for the indicated time. Flies were dissected and anterior and posterior regions of midguts were separated and lysed in a hypotonic buffer (25 mM HEPES, pH 7.5, 5 mM MgCl_2_, 5 mM EDTA, 5 mM DTT, 2 mM PMSF, 10 µg/mL leupeptin, 10 µg/mL pepstatin A) on ice. Midguts were crushed one minute at 30 Hz with the Tissue Lyser. Homogenates were centrifuged at 20 000 g for 45 min at 4°C. The supernatant was the soluble fraction (considered as cytosol-enriched fraction) and the pellet was the insoluble fraction (containing membranes).

Proteins were dosed by Bradford method (Protein Assay Dye Reagent Concentrate, Biorad #500-0006). 20µg of midgut proteins and 2.10^4^ CFU of spores were deposited on a 8.5% SDS-polyacrylamide gel. After migration at 100V for 1h30, proteins were transferred onto a PVDF membrane (Immobilon®-P Membrane, Millipore #IPVH00010) (120mA/gel, 1h) in a semi-dried transfer system with a transfer buffer (200 mM glycine, 25 mM Tris base, 0,1% SDS, pH7,4, 20% methanol). Membranes were saturated with 5% milk in TBS-T (140mM NaCl, 10 mM Tris base, 0,1% Tween 20, pH 7.4) for 1h and incubated overnight at 4°C with a homemade anti-Cry1A antibody (Babin et al., 2020) diluted at 1/7 500 in TBS-T 3% BSA. After three rinsings of 10 min each with TBST, membranes were incubated with an anti-rabbit antibody coupled with HRP (Goat Anti-Rabbit (IgG), Invitrogen #G21234) diluted 1/7 500 in TBS-T 2% milk for 1 hour at room temperature. Membranes were rinsed three times with TBST and once with TBS. Western blots were revealed with enhanced chemiluminescence (luminol and hydrogen peroxide, homemade) on an autoradiographic film (Amersham Hyperfilm™ ECL, GE Healthcare #28906837).

### Measurement, counting and statistical analysis

In all the data presented, the pictures and counting were always performed in the posterior part of the R4 region (http://flygut.epfl.ch/) named R4bc in the flygut site (see also (Buchon et al., 2013; Marianes and Spradling, 2013)). Experiments were independently repeated at least three times. Cells were counted in a fixed area (20.000 μm²) within the R4bc region. Effects of treatments were analyzed using a Kyplot (KP test). Differences were considered significant when P<0.05 (*P≤0.05, **P≤0.01, ***P≤0.001).

## Acknowledgments

We are grateful to all members of the BES team for fruitful discussions. We want to thank Arnaud Felten (GVB Unit, Anses Ploufragan) and Pierre-Emmanuel Douarre (SEL unit, Anses Maisons-Alfort) for their help regarding the WGS analysis of the mutant *Btk*^*Cry1Ab*^. We also thank Laurent Ruel for providing pWA-Gal4. We are grateful to Hiroki Oda for providing the pUAST-DEFL. We thank Olivier Pierre from the microscopy platform of the Sophia Agrobiotech Institute for his help. Our thanks to the Université Côte d’Azur Office of International Scientific Visibility for English language editing of the manuscript.We also thank the Space, Environment, Risk and Resilience Academy of Université Côte d’Azur for their financial support.

## Author contributions

RJ: Conceptualization, Data curation, Formal analysis, Investigation, Visualization, Writing – original draft

RL: Conceptualization, Investigation, Visualization NZP: Formal Analysis, Investigation, Visualization MPNE: Methodology

AF: Formal Analysis, Data curation

RR: Funding acquisition, Validation, Writing – review & editing MB: Methodology, Resources, Data curation,

DO: Conceptualization, Funding acquisition, Validation, Writing – review & editing

AG: Conceptualization, Data curation, Formal Analysis, Data curation, Funding acquisition, Project administration, Supervision, Validation, Visualization, Writing – review & editing

## Funding

This work has been supported by the French government, through the UCAJEDI Investments in the Future project managed by the National Research Agency (ANR) with the reference number ANR-15-IDEX-01 and through the ANR-13-CESA-0003-01, by the Région Provence Alpes Côte d’Azur, by the Département des Alpes-Maritimes, by the Institut Olga Triballat (PR2016-19) and by the ANSES PNR-EST & ECOPHYTO II (EST-2017-2021). RJ was funded by the association AZM & SAADE (Lebanon) and Université Côte d’Azur (ATER). RL was funded by the Ministère de l’Education Nationale, de l’Enseignement Superieur et de la Recherche (MESR) and a grant from the Fondation pour la Recherche Médicale (FRM).

## Notes

### Competing Interest Statement

The authors have declared no competing interest.

