## Supplemental figures for "*Bacillus thuringiensis* Cry1A toxins divert progenitor cells toward enteroendocrine fate by decreasing cell adhesion with intestinal stem cells"

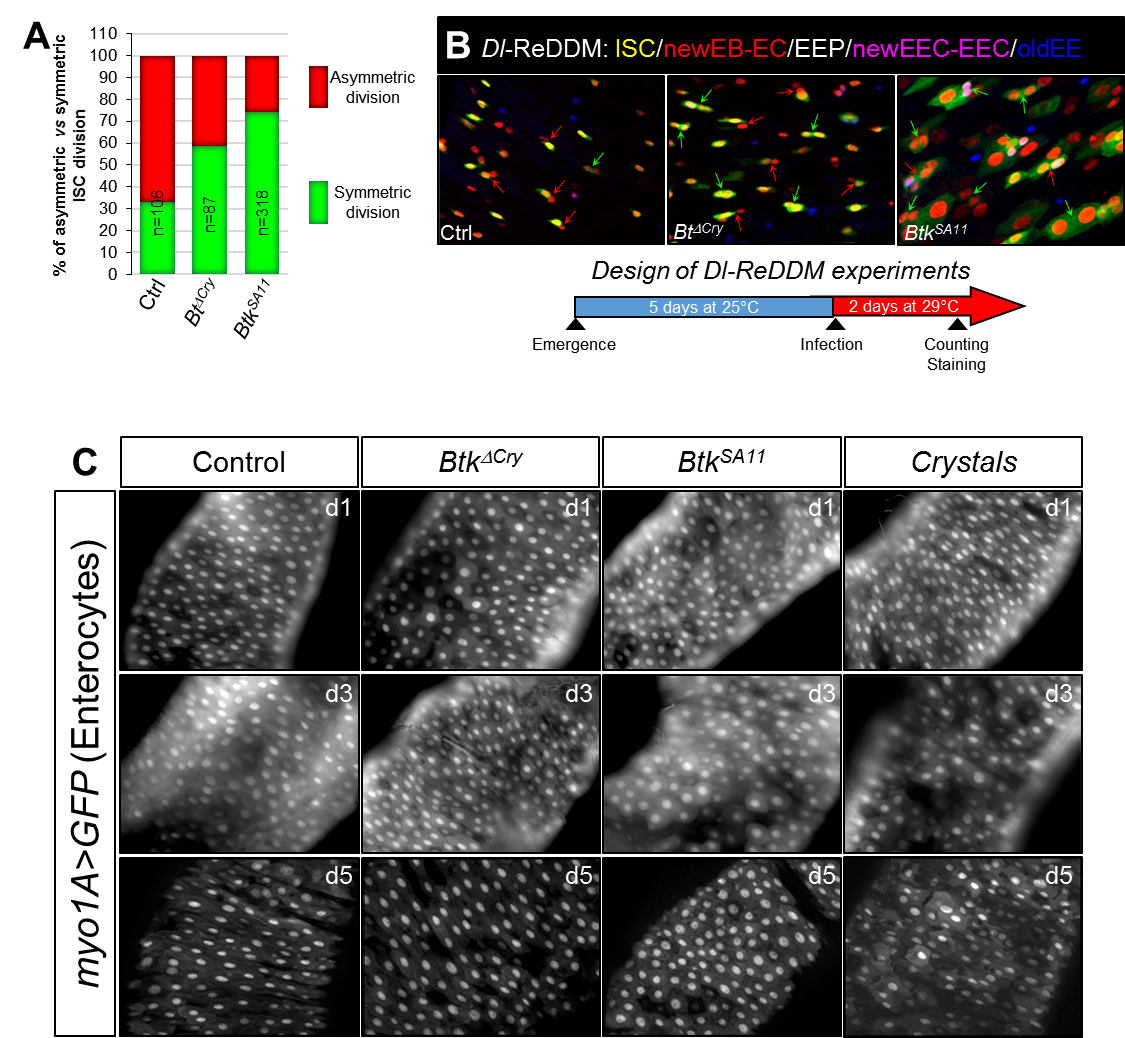

**Figure S1: Crystals of *Btk* Cry protoxins induce symmetric division of ISCs**

(**A** and **B**) Monitoring of the ISC mode of division in posterior midguts of *Dl-ReDDM* flies. *Drosophila* were fed with water, *Btk^∆Cry^* spores or *Btk^SA11^* spores. The experimental design is shown below the panels in (B). In (A) red bars correspond to asymmetric divisions and green bars to symmetric divisions. In (B) anti-Pros (blue) marks the EEPs and EECs. Red arrows point to asymmetric divisions and green arrows point to symmetric divisions.

(**C**) *myo1A>GFP* posterior midguts of *Drosophila* fed with water (control), *Btk^∆Cry^* spores, *Btk^SA11^* spores or crystals at 1, 3 and 5 days PI. GFP marks the ECs.

n=number of samples.

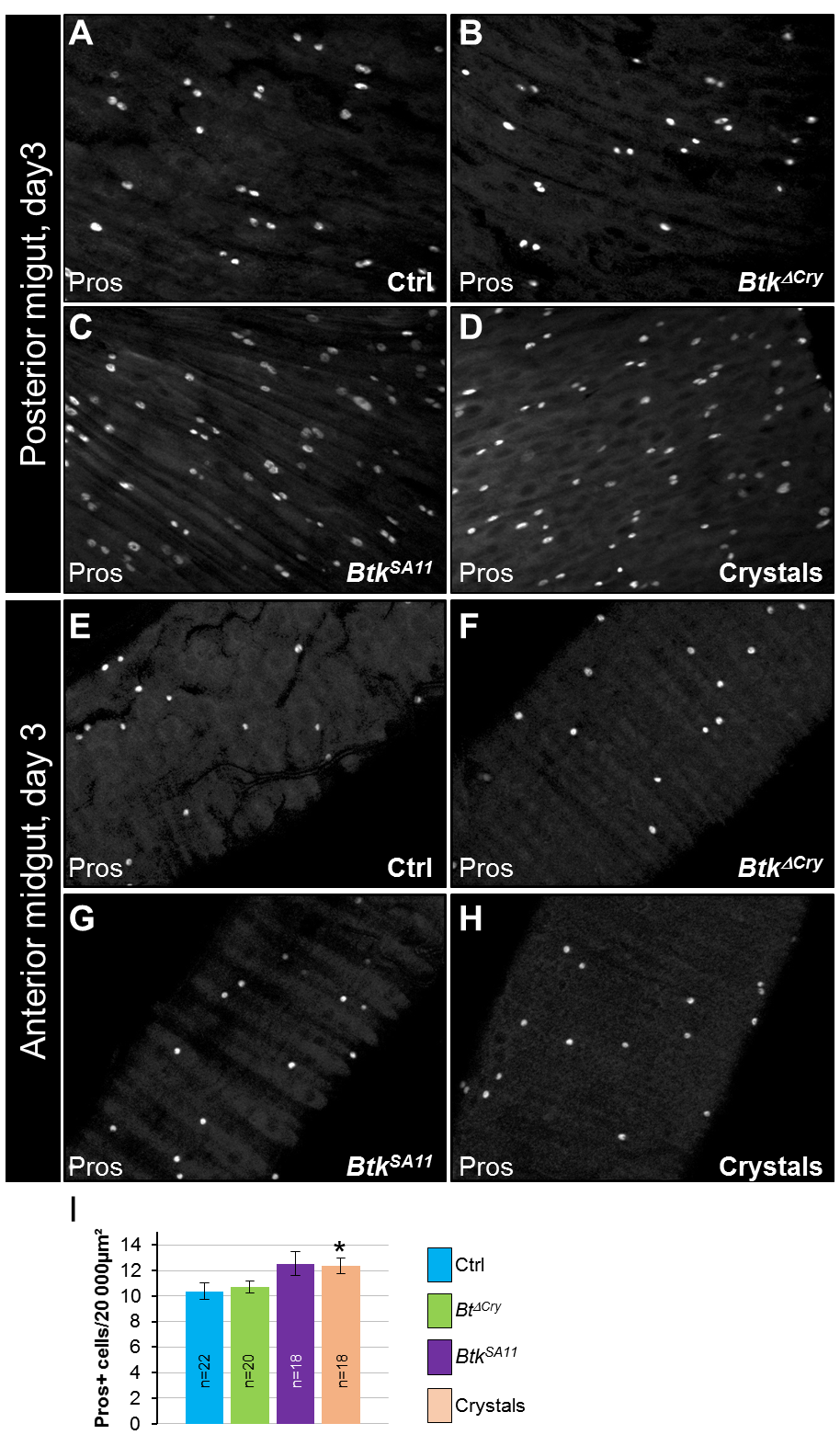

**Figure S2: *Btk^SA11^* crystals induce an increase EEP and EEC number in the posterior midgut**

(**A-H**) Anti-Pros was used to mark the EECs and EEPs

(**A-D**) R4 region of *esg>GFP* midguts 72h PI of water (A), *Btk^∆Cry^* spores (B), *Btk^SA11^* spores (C) or crystals (D).

(**E-H**) R2 of region *esg>GFP* midguts 72h PI of water (E), *Btk^∆Cry^* spores (F), *Btk^SA11^* spores (G) or crystals (H)

(**I**) Number of Pros+ cells in the anterior part of the midgut 72h PI of water (blue), *Btk^∆Cry^* spores (green), *Btk^SA11^* spores (purple) or Crystals (pink). n=number of samples; * (P ≤ 0.05).

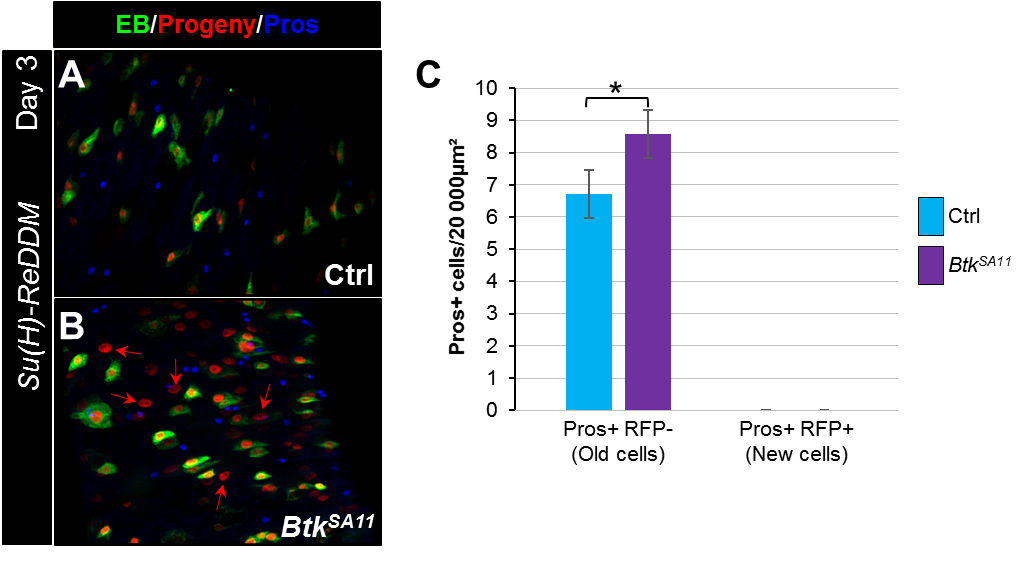

**Figure S3: EBs do not give birth to EECs**

(**A** and **B**) R4 region of *Su(H)- ReDDM* flies. Flies were fed with either water (A, Ctrl) or *Btk^SA11^* spores (B). Midguts were labelled for Pros (blue).

(**C**) Counting old EECs (Pros+ RFP-) and new EECs (Pros+ RFP+) in the conditions described in (A and B). n=number of samples; ns=not significant; * (P ≤ 0.05).

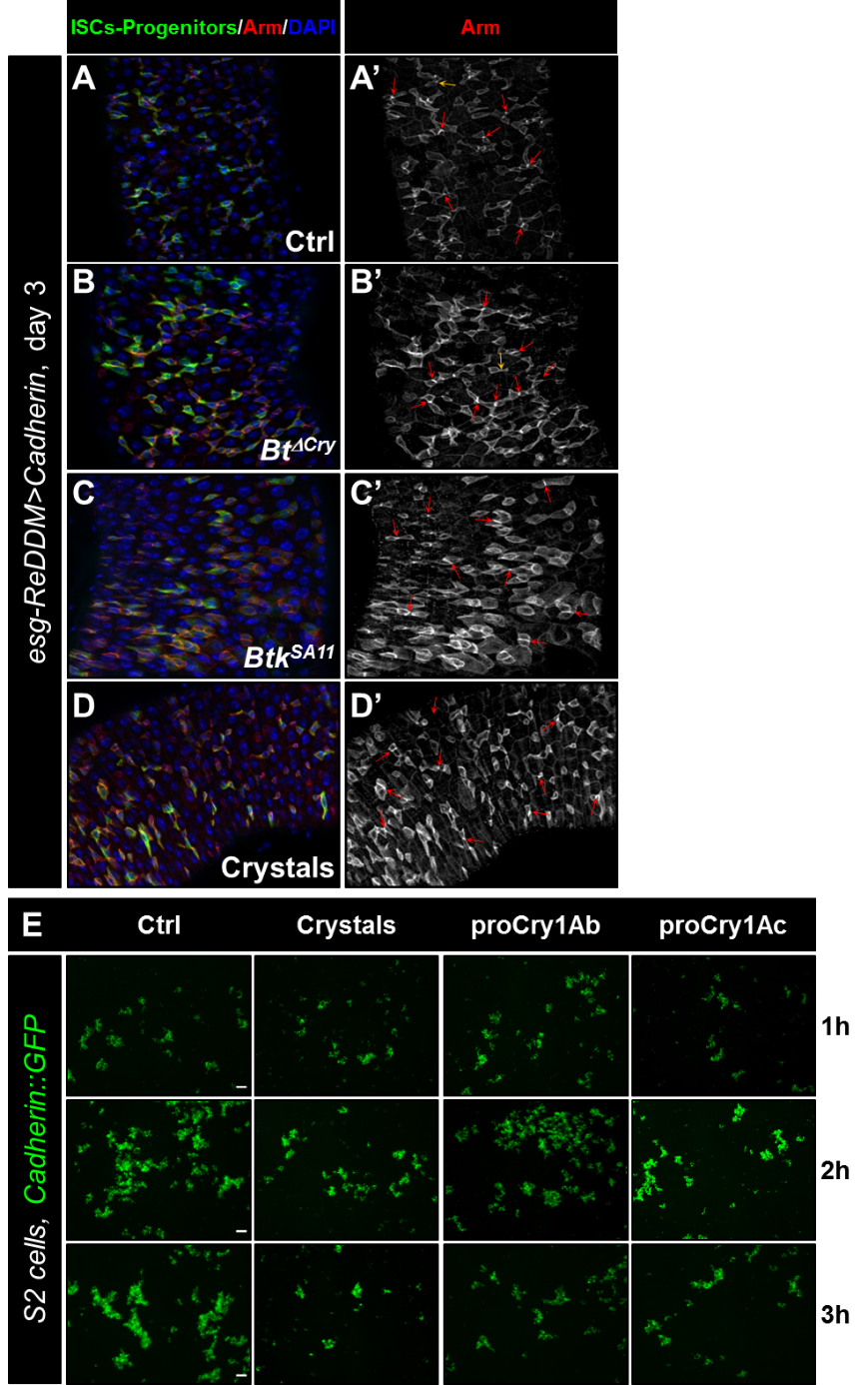

**Figure S4: Cry1A protoxins reduced homophilic interactions of DE-cadherin**

(**A-D’**) *esg-ReDDM>DE-Cad Drosophila* midgut R4 region. Flies were fed with water (A and A', Ctrl), *Btk^∆Cry^* spores (B and B’), *Btk^SA11^* spores (C and C’) or Crystals (D and D') and observed 72h PI (see Figure 3A for the experimental design). Midguts were labeled with anti-Arm (red) which strongly marks the adherent junctions between ISC and progenitors (green). In blue Dapi marks the nuclei. Red arrows point to the high intensity of adherens junction staining. Yellow arrows point to the weak intensity of adherens junction staining.

(**E**) Representative images of cell aggregation assays obtained from 3 independent experiments.

S2 cells transiently expressing the DE-Cadherin::GFP were placed under constant rotation and incubated for 1, 2 or 3h with or without *Btk* crystals or purified Cry1Ab or Cry1Ac protoxins. Scale bars represent 50 µm.

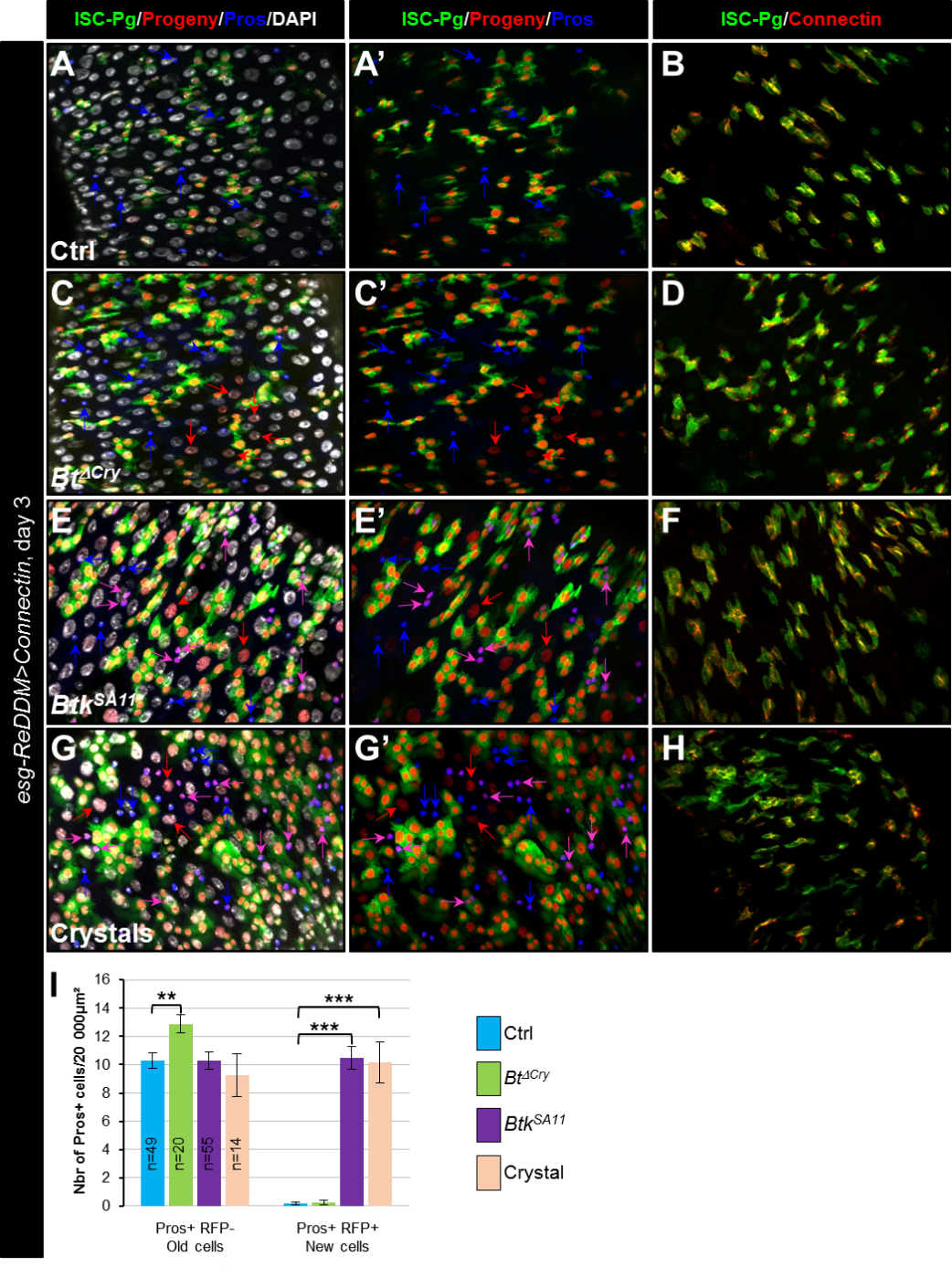

**Figure S5: Connectin overexpression does not rescue cell adhesion disturbance induced by *Btk* crystals of toxins**

(**A-I**) *esg-ReDDM>connectin*. These flies specifically overexpress Connectin in ISCs and progenitors. Flies were fed with water (A-A’, B and blue in I, Ctrl), *Btk^∆Cry^* spores (C, C’, D and green in I), *Btk^SA11^* spores (E, E’, F and purple in I) or Crystals (G-G’, H and pink in I) and observed 72h PI. Midguts were stained for Pros (blue in A, A', C, C', E, E', G and G') and for Connectin (red in B, D, F, and H). DAPI marks the nuclei (white in A; C, E and G). Blue arrows point to old EECs, pink arrows point to new EECs and red arrows newborn ECs.

(**I**) Number of old cells (Pros+ RFP-) and new cells (Pros+ RFP+) in the conditions described in (A-H).

n=number of samples, ** (P ≤ 0.01), *** (P ≤ 0.001).

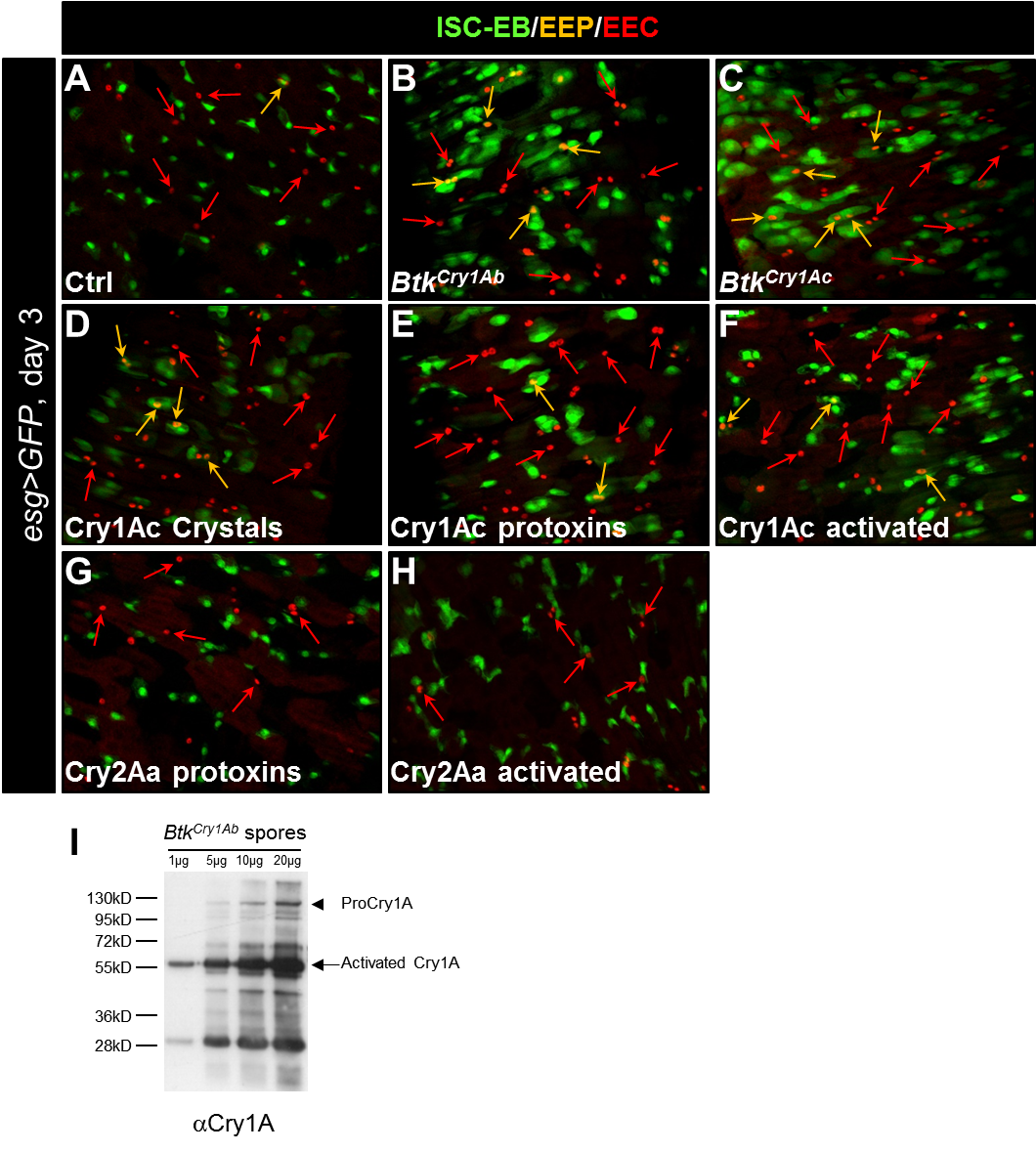

**Figure S6: Cry1A toxins mimic *Btk* crystal effects**

(**A-H**) esg>GFP *Drosophila* midgut R4 region 72h PI of water (A, Ctrl), *Btk*^Cry1Ab^ spores, (B), *Btk^Cry1Ac^* spores (C), Cry1Ac crystals (D), Cry1Ac protoxins (E), Cry1Ac activated toxins (F), Cry2Aa protoxins (G) and Cry2Aa activated toxins (H). GFP labels ISCs and progenitors, Pros labels EEPs and EECs (red). Orange arrows point to EEPs and red arrows points to the EECs.

(**I**) Western blot on *Btk^Cry1Ab^* spores using the Anti-Cry1A polyclonal antibody. Spores of *Btk^Cry1Ab^* were resuspended in water to obtain a concentration of 1µg/µL. 1, 5, 10 and 20 µg of proteins were loaded on a 10% gel of acrylamide. The protoxins (130kDa, arrowhead) are present but in very low quantities. The activated 67kDa forms (arrow) are more abundant.
