## supplemental information for "*Bacillus thuringiensis* Cry1A toxins divert progenitor cells toward enteroendocrine fate by decreasing cell adhesion with intestinal stem cells"

**Reagents and Tools table:**

| Reagent /ressource | Reference or source | Identifier or catalog number |
| --- | --- | --- |
| **Bacterial strains** |  |  |
| B*. thuringiensis*: *Btk* SA11 | Bioinsecticide | Delfin |
| *B. thuringiensis*: *Btk^∆Cry^* | BGSC | #4D22 |
| *B.thuringiensis*: *Btk* producing Cry1Ac | BGSC | #4D4 |
| *B. thuringiensis*: *Btk* producing Cry1Ab | This study |  |
| *E. Coli* producing Cry1Ab | BGSC | #ECE54 |
| *E. Coli* producing Cry1Ac | BGSC | #ECE53 |
| *E. Coli* producing Cry2Aa | BGSC | #ECE126 |
| **Cell Culture** | | |
| *Drosophila melanogaster* Schneider 2 (S2) cells | Gift from L. Ruel | (Gallet et al., 2006) |
| Schneider’s insect medium | Sigma-Aldrich | Cat# S0146 |
| TransIT®-2020 | Mirus Bio | Cat# MIR5400 |
| **Recominant DNA** | | |
| pUAST-DECadherin tagged with GFP (DEFL) |  | (Oda and Tsukita, 1999) |
| pWA-Gal4 | Gift from L. Ruel |  |
| **Antibodies** | | |

| Mouse anti-Armadillo (ß-catenin) antibody | DSHB | Cat# N27A1  RRID:AB_528089 |
| --- | --- | --- |
| Mouse anti-Connectin | DSHB | Cat# Connectin C1.427, RRID:AB_1066083 |
| Mouse anti-Prospero antibody | DSHB | Cat# MR1A  RRID:AB_528440 |
| Rabbit anti-Cleaved Caspase-3 (Asp175) | Cell Signalling | Cat# 9661  RRID:AB_2341188 |
| Rabbit anti-phospho-Histone H3 (Ser10) | Millipore | Cat# 06-570  RRID:AB_31017 |
| Rabbit polyclonal anti-Cry1A |  | (Babin et al., 2020) |
| Goat anti mouse IgG (H+L) secondary antibody, AlexaFluor-647 | Invitrogen | Molecular Probes Cat# A-21235, RRID:AB_2535804 |
| Goat anti mouse IgG (H+L) secondary antibody, AlexaFluor-546 | Invitrogen | Molecular Probes Cat# A-11003, RRID:AB_141370 |
| Goat anti-rabbit IgG (H+L) secondary antibody, AlexaFluor-647 | Invitrogen | Thermo Fisher Scientific Cat# A32733, RRID:AB_2633282 |
| Goat anti-rabbit IgG (H+L) secondary antibody, AlexaFluor-546 | Invitrogen | Thermo Fisher Scientific Cat# A-11010, RRID:AB_253407 |
| **Other reagents** | | |
| PBS 10x | Euromedex | ET330 |
| Formaldehyde 16% | Thermo scientific | Cat# 28908 |
| Fluoroshield-DAPI | Sigma | F6057-20mL |

| ***Drosophila* strains** |  |  |
| --- | --- | --- |
| WT canton S | <https://bdsc.indiana.edu/> | #64349 |
| *w; Sco/CyO; tub-GAL80^ts^/TM6b* | <https://bdsc.indiana.edu/>  M. Vidal | #7018 |
| *w; tub-GAL80^ts^; TM2/TM6b* | <https://bdsc.indiana.edu/> | #7019 |
| *w; ; Dl-GAL4/TM6b* | S. Hou and X. Zeng | (Zeng et al., 2010) |
| *w; tub-GAL80^ts^; Dl-GAL4 UAS-GFP/TM6b* | This study |  |
| *w; esg-GAL4^NP5130^* | <https://bdsc.indiana.edu/>  N. Tapon | #67054 |
| *w; esg-GAL4^NP5130^ UAS-GFP* | N. Tapon | (Shaw et al., 2010) |
| *w; esg-GAL4^NP5130^ UAS-GFP; tubGAL80^ts^* | Y. Apidianakis | (Apidianakis et al., 2009) |
| *w; Su(H)GBE-GAL4, UAS-CD8::GFP* | M. Vidal | (Zeng et al., 2010) |
| *w; Su(H)GBE-GAL4/SM6β; tub-GAL80^ts^ UAS-GFP/TM6b* | This study |  |
| *w; myo1A-GAL4* | N. Tapon | (Shaw et al., 2010) |
| *w; myo1A-GAL4 UAS-GFP/CyO* | Y. Apidianakis | (Apidianakis et al., 2009) |
| *w; UAS-GFP/TM3 Sb* | <https://bdsc.indiana.edu/> | #5430 |
| *w; UAS-GFP::CD8; UAS-H2B::RFP/TM2* | T. Reiff and M. Dominguez | (Antonello et al., 2015) |
| *w; UAS-CD8::GFP; UAS-H2B::RFP, tub-GAL80^ts^/TM2* | T. Reiff and M. Dominguez | (Antonello et al., 2015) |
| *w; esg-GAL4, UAS-CD8::GFP/CyO; UAS-H2B::RFP, tub-GAL80ts/TM6b* | T. Reiff and M. Dominguez | (Antonello et al., 2015) |
| *w; UAS-CD8::GFP; Dl-GAL4, UAS-H2B::RFP/TM6b* | This study |  |
| *w; ; UAS-GC3Ai^G7S^ (UAS-Casp::GFP)* | M. Suzanne | (Schott et al., 2017) |
| *w; UAS-shg-R (DE-Cadherin)* | <https://bdsc.indiana.edu/> | #58494 |
| *w; UAS-connectin* | JP Boquete and B. Lemaitre | (Zhai et al., 2017) |
| *y w, shg::Tomato* | <https://bdsc.indiana.edu/> | #58789. |
| ***Dl-ReDDM*** *(w/w; UAS-CD8::GFP/UAS-CD8::GFP; Dl-GAL4, UAS-H2B::RFP/UAS-H2B::RFP, tub-GAL80^ts^)* | This study |  |
| ***esg-ReDDM*** *(w/w^+^; esg-GAL4, UAS-CD8::GFP/+; UAS-H2B::RFP, tub-GAL80^ts^/+)* | This study |  |
| ***Su(H)-ReDDM*** *(w/w; Su(H)-GAL4/UAS-GFP::CD8; tub-GAL80^ts^ UAS-GFP/UAS-H2B:RFP)* | This study |  |

Antonello, Z. A., Reiff, T., Ballesta-Illan, E., and Dominguez, M. (2015). Robust intestinal homeostasis relies on cellular plasticity in enteroblasts mediated by miR-8-Escargot switch. EMBO *34*, 2025-2041. doi: 2010.15252/embj.201591517. Epub 201592015 Jun 201591515.

Apidianakis, Y., Pitsouli, C., Perrimon, N., and Rahme, L. (2009). Synergy between bacterial infection and genetic predisposition in intestinal dysplasia. Proc Natl Acad Sci U S A *106*, 20883-20888. Epub 22009 Nov 20823.

Babin, A., Nawrot-Esposito, M.-P., Gallet, A., Gatti, J.-L., and Poirié, M. (2020). Differential side-effects of Bacillus thuringiensis bioinsecticide on non-target Drosophila flies. Scientific Reports *10*, 16241.

Gallet, A., Ruel, L., Staccini-Lavenant, L., and Therond, P. P. (2006). Cholesterol modification is necessary for controlled planar long-range activity of Hedgehog in Drosophila epithelia. Development *133*, 407-418. Epub 2006 Jan 2005.

Oda, H., and Tsukita, S. (1999). Nonchordate classic cadherins have a structurally and functionally unique domain that is absent from chordate classic cadherins. Dev Biol *216*, 406-422.

Schott, S., Ambrosini, A., Barbaste, A., Benassayag, C., Gracia, M., Proag, A., Rayer, M., Monier, B., and Suzanne, M. (2017). A fluorescent toolkit for spatiotemporal tracking of apoptotic cells in living Drosophila tissues. Development *144*, 3840-3846. doi: 3810.1242/dev.149807. Epub 142017 Sep 149804.

Shaw, R. L., Kohlmaier, A., Polesello, C., Veelken, C., Edgar, B. A., and Tapon, N. (2010). The Hippo pathway regulates intestinal stem cell proliferation during Drosophila adult midgut regeneration. Development *137*, 4147-4158. doi: 4110.1242/dev.052506. Epub 052010 Nov 052510.

Zeng, X., Chauhan, C., and Hou, S. X. (2010). Characterization of midgut stem cell- and enteroblast-specific Gal4 lines in drosophila. Genesis *48*, 607-611. doi: 610.1002/dvg.20661.

Zhai, Z., Boquete, J. P., and Lemaitre, B. (2017). A genetic framework controlling the differentiation of intestinal stem cells during regeneration in Drosophila. PLoS Genet *13*, e1006854. doi: 1006810.1001371/journal.pgen.1006854. eCollection 1002017 Jun.
